# Induced Cognitive Impairments Reversed by Grafts of Neural Precursors: a Longitudinal Study in a Macaque Model of Parkinson’s Disease

**DOI:** 10.1101/2020.12.17.423293

**Authors:** Florence Wianny, Kwamivi Dzahini, Karim Fifel, Charles R.E. Wilson, Agnieszka Bernat, Virginie Dolmazon, Pierre Misery, Camille Lamy, Howard M. Cooper, Emmanuel Procyk, Henry Kennedy, Pierre Savatier, Colette Dehay, Julien Vezoli

**Author notes:** Corresponding author. (J.V.); (C.D.). International Institute for Integrative Sleep Medicine (WPI-IIIS), University of Tsukuba; Tsukuba, Ibaraki 305-8575, Japan. Laboratory of Molecular Diagnostics, Department of Biotechnology, Inter-collegiate Faculty of Biotechnology, University of Gdańsk and Medical University of Gdańsk; Gdańsk, Poland (Agnieszka Bernat). Laboratory of Experimental Embryology, Institute of Genetics and Animal Biotechnology, Polish Academy of Sciences; Warsaw, Poland (Agnieszka Bernat).

## Abstract

Parkinson’s disease (PD) evolves over an extended and variable period in humans; several years prior to the onset of classical motor symptoms, cognitive deficits as well as sleep and biological rhythm disorders develop and worsen with disease progression, significantly impacting the quality of life of patients. The gold standard MPTP macaque model of PD recapitulates the progression of motor and non-motor symptoms over contracted periods of time.

Here, this multidisciplinary and multiparametric study follows, in five animals, the steady progression of motor and non-motor symptoms and describes their reversal following bilateral grafts of neural precursors in diverse functional domains of the basal ganglia.

Results show unprecedented recovery from cognitive symptoms in addition to a strong clinical motor recuperation. Both motor and cognitive recovery and partial circadian rhythm recovery correlate with the degree of graft integration into the host environment as well as with in-vivo levels of striatal dopaminergic innervation and function.

Given inter-individuality of disease progression and recovery the present study underlines the importance of longitudinal multidisciplinary assessments in view of clinical translation and provides empirical evidence that integration of neural precursors following transplantation efficiently restores function at multiple levels in parkinsonian non-human primates.

**One Sentence Summary:** Empirical evidence that cell therapy efficiently reverts cognitive and clinical motor symptoms in the non-human primate model of Parkinson’s disease.

## INTRODUCTION

Parkinson’s disease (PD) is a neurodegenerative condition affecting up to 10M of the worldwide population with incidences increasing with age, making PD the fastest growing neurological disorder in a globally aging population [1]. The clinical manifestation is primarily motor with a typical parkinsonian syndrome including bradykinesia, rigidity and resting tremor and is confirmed by clinical diagnostic tools including *in-vivo* imaging of the denervation of the nigrostriatal dopaminergic (DA) axis [2]. Unfortunately, 60-80% of DA cells are lost prior to the onset of clinically diagnosed motor symptoms [3]. During this so-called premotor period preceding the clinical threshold, the DA-lesion is progressive and accompanied by the manifestation and further deterioration of non-motor symptoms including cognitive disorders as well as perturbation of circadian rhythm and sleep disorders [4, 5]. Although considered secondary, non-motor symptoms nonetheless have a significant impact on quality of life; the cognitive abilities of the vast majority of PD patients deteriorate leading to psychiatric disturbances [6] with circadian perturbations affecting biological rhythms [7].

DA lesion is the hallmark of PD and in addition to circadian regulation affects multiple frontal lobe functions including performance monitoring, motivational aspects of behavior and motricity. However, there is currently no cure for PD and palliative therapies such as levodopa and Deep-Brain Stimulation (DBS), mainly correct motor symptoms and often give rise to behavioral perturbation [8]. For example, DBS shows no long-lasting effect on axial symptoms e.g. freezing of gait and negatively impacts cognitive and neuropsychiatric symptoms [9, 10]. Importantly palliative therapies do not solve the issue of neuronal loss and the long-term prognosis. Consequently, cell replacement therapy for PD has been recently re-evaluated as a potential cure for PD [11], leading to recent trials [12] aimed at improved grafting procedures, and to pave the way for stem-cell based transplantation in humans. In this respect there is a recognized need for more detailed perspectives from pre-clinical investigations in animal models [13].

With the aim of optimizing the therapeutic use of cell replacement in PD we have examined the global consequence of cell grafts at different stages of the disease in a macaque model of PD. While numerous efforts have been deployed to use non-motor symptoms as premotor markers for the prognosis of PD [14], virtually no studies have addressed the impact of cell replacement on non-motor symptoms [15] leading to insufficient appraisal of their multi-parametric impact at either pre-clinical or clinical stages [8, 16]. This is problematic because advanced PD with an extensive nigrostriatal DA lesion constitutes a highly deleterious environment for the survival of grafted cells [17, 18]. Hence in the present study we use an experimental design that allows us to address the capacity of cell grafts to reverse the disease at disease onset in the motor phase and assess the impact of DA lesion extent on graft efficiency. This design also allowed us to characterize the dynamics of graft induced recovery by comparing it to spontaneous recovery in the premotor stage. Because non-motor symptoms have a gradual onset prior to clinical manifestations of DA denervation, evaluation of the effects of NP grafts in parkinsonian non-human primates (NHP) requires a longitudinal multi-parametric investigation, which we have implemented in the present study. We grafted neural precursors (NPs) because they hold the promise of enhanced efficiency compared to DA neurons [19–21], as in addition to their neuro-restorative potential NPs have the capacity to differentiate into neuroprotective astroglial cells [22–24].

Repeated systemic injections of low-doses of MPTP (0.2 mg/kg every 3-4 days) induced a parkinsonian syndrome with a slow and typically dorso-ventral progression of the nigrostriatal denervation, together with a premotor expression of cognitive and circadian *i.e.* non-motor symptoms [25–27]. Here, we implemented a multidisciplinary / multiparametric approach to assess the therapeutic potential of bilateral grafts of NPs in multiple functional domains of the basal ganglia of NHP (*macaca fascicularis*). For ethical reasons we reduced the number of animals used by monitoring individual macaques so that each case constitutes its own control. This required continuous assessment of clinical motor and non-motor symptoms as well as *in-vivo* monitoring of the nigrostriatal lesion with Positron Emission Topography (PET) imaging of DA transporters (DAT). Monitoring was prior to and following the induction of stable motor symptoms [27], and subsequent to transplantation of NPs. In addition, Fluoro-DOPA (^18^F-DOPA) imaging allowed assessment of the impact of the NP grafts on DA function [28]. Finally, *post-mortem* immunohistological examination of NP grafts was performed in order to evaluate the impact of the graft on host tissue as well the survival, integration and differentiation fate of the grafted cells into the host brain.

## RESULTS

The experimental design (**Fig.1A, Table S1**) allowed full characterization of MPTP intoxication. In Stage I, all animals show non-motor symptoms [25, 26] and a nigrostriatal DA denervation superior to 70% [27] but, at this stage, spontaneous recovery from motor symptoms occurs and this was investigated in 3 out of the 6 individual cases [25, 27, 29]. Comparison of cognitive, circadian and DA function markers enabled characterization of spontaneous and graft-induced recovery. Subsequent to Stage I, resuming MPTP injections until a clinical score of 10 is reached on two consecutive days [27] leads to a Stage II, where motor symptoms become permanent. In order to implement this experimental strategy it is necessary to adapt cumulated MPTP-doses to individual clinical motor scores (**Table S2**) thereby taking into account the known variability of response to MPTP and adjustment of the intoxication protocol (**Fig.1A**, **Table S2**, see **Materials and Methods**). Clinical scores and behavioral measures were divided into quantiles (Q1-5, as described in [25], see **Materials and Methods**) in order to compare cases on the basis of the presumed DA-lesion rather than on the time spent in the premotor (MPTP), motor (post-MPTP/pre-graft) and post-graft periods. We compared groups (*recovered* vs. *non-recovered*) across control, premotor and motor quantiles (pre-/post-MPTP) to evaluate differences in behavioral markers between groups due to MPTP intoxication and, across pre- and post-grafts quantiles (pre-/post-graft) in order to evaluate differences between groups due to NP grafts (one-way anova: F-test (groups,df), p-value, effect-size 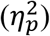, see **Materials and Methods**).

**Fig. 1.**
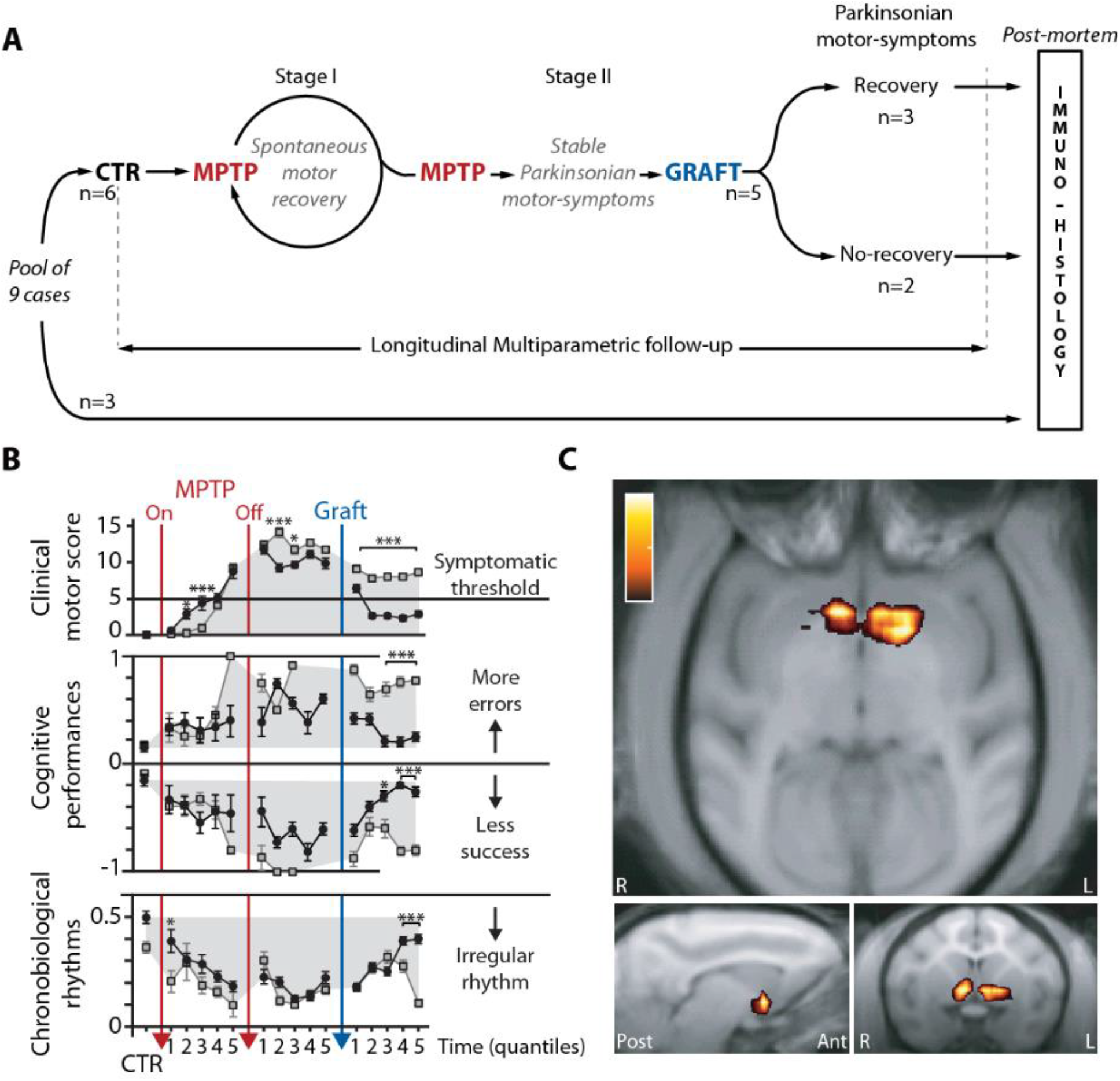
Motor, non-Motor and Functional Markers of Graft-induced Recovery. (**A**) Study design. At early stages of MPTP treatment (Stage I) there is spontaneous recovery from motor symptoms but not from cognitive and circadian symptoms. Continued MPTP intoxication leads to Stage II where motor symptoms now become permanent. (**B**) Cases are grouped in *recovered* (black) vs. *non-recovered* (grey) according to the level of parkinsonian motor-symptoms following cell therapy. Case2 excluded because of per-operative brain hemorrhage following graft. Motor symptoms clinically scored with the Parkinsonian Monkey Rating Scale (PMRS top panel), Cognitive performances (range normalized to control performance, −1 for success, middle panel) and, Chronobiological rhythms reflected by the ratio of spontaneous locomotor activity during the light and dark periods (normalized contrast with Light/Dark ratio in control conditions, bottom panel). Means ± SE, anova between groups, adjusted p-values for multiple comparisons * < 0.05, *** < 0.001, see **Table S8**. (**C**) Increase of nigrostriatal innervation. Parametric maps of DAT binding potential with axial (top), sagittal (bottom-left panel) and coronal (bottom-right panel) views of significant positive difference for the contrast Pre-vs. Post-Graft (color scale from 0.2-1, case1, recovered group). See **Fig.S1** for individual behavioral markers and **Fig.S9** for statistical thresholding in (**C**) and negative difference.

### Behavioral and functional markers of chronic MPTP-induced Parkinsonism

During Stage II, all cases displayed a stable parkinsonian syndrome with clinical motor scores ranging from mild to severely symptomatic (**Figs.1B, S1A**). In two out of the five grafted cases, we failed to observe recovery from motor symptoms whereas the three other cases exhibited robust recovery from motor symptoms with a clinical motor score returning below the symptomatic threshold (pre-/post-graft: *F*(2,10) = 126, *p* < 2*E*-16, 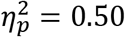 while effect-size was clearly reduced over pre-/post-MPTP quantiles: *F*(2,11) = 11, *p* < 2*E*-16, 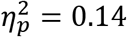, **Figs.1B, S1A, Table S8**). We accordingly assigned cases to the *non-recovered* or *recovered* group based on clinical motor outcome following NP grafts (**Figs.1A-B, S1A**).

During the pre-Graft period, non-motor symptoms emerged in all animals with progressive deterioration during the premotor period *i.e*. before motor scores reached clinical thresholds (**Figs.1B, S1**). Alterations of circadian rhythms appeared progressively during the premotor phase (**Figs.1B, S1C**) to reach peak levels during the symptomatic period as previously described [26]. This pattern of decreased (increased) activity during the Light (Dark) phase resembles circadian alterations of the sleep-wake cycle observed in PD patients which manifests as increased daytime sleepiness and fragmented sleep structure [30]. The premotor and motor alteration of rest-wake locomotor rhythms in all cases exhibited reduced light/dark (LD) ratios (minimal effect-size of group difference across pre-/post-MPTP quantiles: *F*(2,11) = 4.3, *p* = 3.2*E*-16, 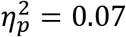, **Fig.1B, Table S8**) and relative amplitude, as well as increased rhythm variability and fragmentation (**Fig.S1C**). In all cases cognitive performance deteriorated during MPTP intoxication and after offset (pre-/post-MPTP quantiles, no group difference in Success: *F*(2,11) = 1.8, *p* > 0.05, 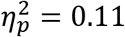, and similarly minor effect-size of group for Errors: *F*(2,11) = 2.1, *p* = 0.03, 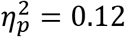, **Figs.1B, S1B**).

Nigrostriatal DA innervation evaluated by striatal DAT binding, decreased by more than 80% relative to control measures of striatal DAT levels with a relatively spared binding in the ventral striatum [27]. This dorsoventral pattern of the nigrostriatal lesion was later confirmed by semi-quantitative evaluation of *post-mortem* immunostaining for DAT and TH (**Fig.2**) with a significantly more advanced lesion in *non-recovered* cases (**Fig.2C**).

**Fig. 2.**
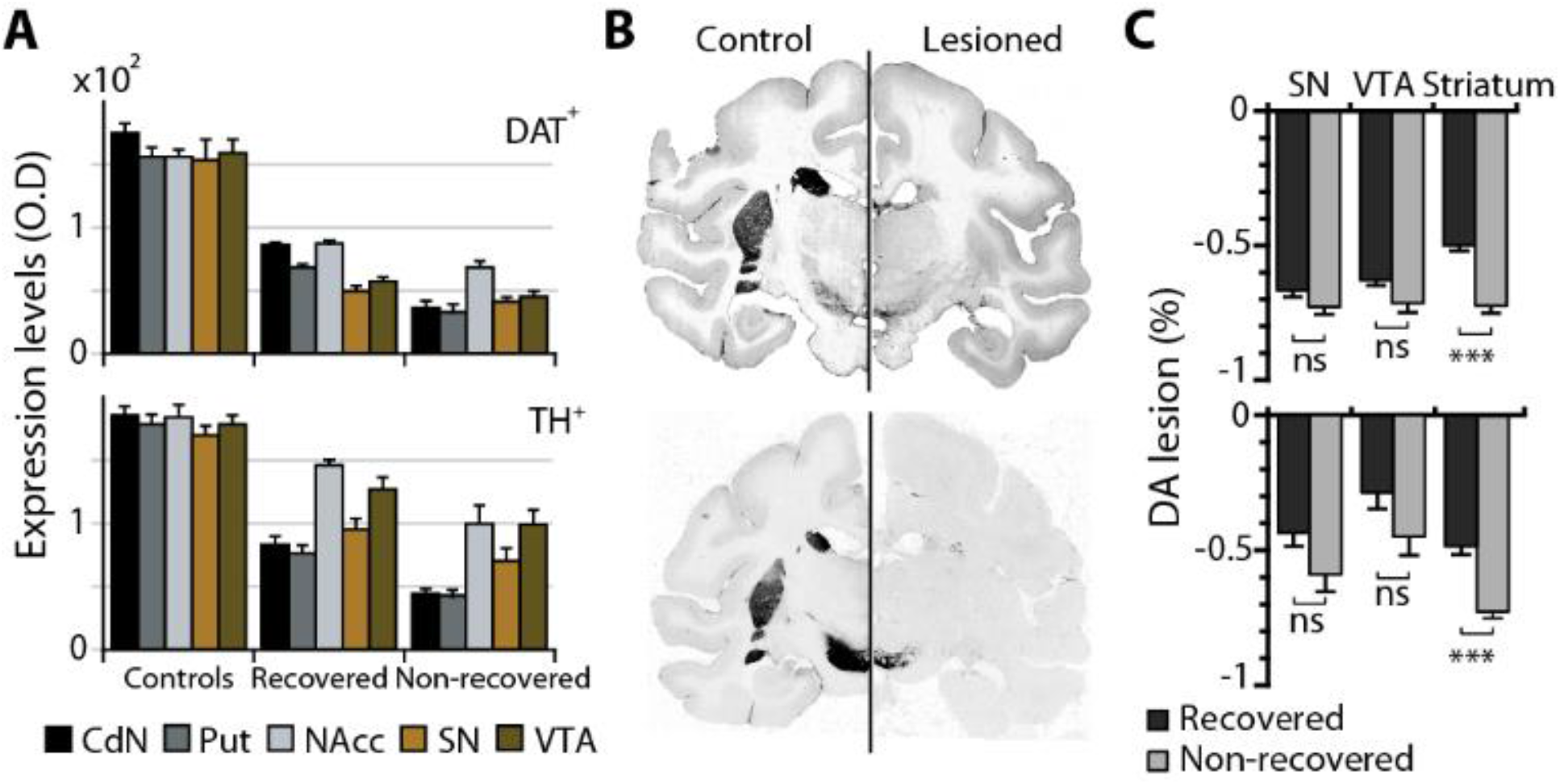
Dopaminergic nigrostriatal lesion. Immunostaining for DAT (upper row) and for TH (lower row). (**A**) Optical density (O.D.) measures of DA lesion (arbitrary units) is made on post-mortem tissue of controls, *recovered* MPTP, *non-recovered* MPTP. (**B**) Illustrative histological sections showing immunostaining. (**C**) Percent change in expression levels between MPTP and Controls provides DA lesion estimates by comparing MPTP sub-groups (two-sided Wilcoxon rank sum test, Bonferroni correction for multiple comparisons *** p < 0.001, Z ≥ 5.75), *Recovered* (black) vs. *Non-recovered* (gray). Mean±SE. *Recovered* group: cases 1, 4 and 5; *Non-recovered* group: cases 3 and 6; Controls group: n=3.

### Behavioral and functional markers following NP grafts

NPs were derived from a rhesus embryonic stem cell line (LYON-ES1) stably expressing the fluorescent marker tau-GFP [31, 32], which allows precise tracking of the grafted NPs and their integration into the host circuits. We chose to graft NPs given their inherent neuroprotective potential resulting from interaction with host-cells [33] and their observed higher survival rate compared to mature DA cell grafts [22, 34, 35]. Because *in-vitro* NPs show both astroglial and neuronal-including DA-fates (**Fig.S2**), it is possible to assess the influence of the host-environment on the differentiation fate of transplanted cells. NP grafting was delayed by 3 months on average following the last MPTP injection (13±6weeks, see **Materials and Methods**). Bilateral transplantations in multiple functional domains of the basal ganglia potentiated graft efficiency on both motor and non-motor symptoms. Grafts were targeted in striatal regions involved in cognitive (*i.e*. anterior caudate nucleus – ant-CdN) and motor functions (*i.e.* posterior putamen – post-Put.), as well as in the substantia nigra (SN), the latter providing the source of nigrostriatal DA innervation *i.e*. *pars compacta*, and part of the frontal DA innervation.

Following NP grafts, the reduction of the clinical motor score was associated with an additional cognitive and circadian recovery in the *recovered* group (**Figs.1B, S1**). In the *recovered* group, clinical scores progressively and significantly diverged at the 2^nd^ quantile from those of the *non-recovered* group, whereas group differences were significant after the 3^rd^ quantile for non-motor symptoms (**Fig.1B**). Clinical scores dropped below symptomatic thresholds on average 6 weeks following NP grafts, to ultimately achieve maximal motor recovery 5 weeks later (**Figs.1B, S1A**). In the *non-recovered* group, the initial cognitive and LD ratio recovery shortly after the graft was followed by a relapse on the last three to two quantiles (**Fig.1B**). However, the effect of the group difference of non-motor markers in pre-/post-graft was 3-5 times higher than in pre-/post-MPTP (with large effect-size for Success: *F*(2,10) = 10, *p* = 3.1*E*-13, 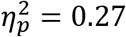 and Errors: *F*(2,10) = 11, *p* = 1.6*E*-14, 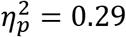, and moderate for LD ratio: *F*(2,10) = 12, *p* < 2*E*-16, 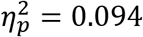, **Fig.1B, Table S8**). Non-parametric circadian rhythm analyses (see **Material and Methods**) revealed high variability across cases, preventing post-graft group comparison on those markers (**Fig.S1C**). In the *recovered* group, markers of striatal DA function and DAT increased following NP grafts, in correlation with the observed behavioral recovery (**Figs.1B-C, 3B-C**). Contrasting with the beneficial effects of NP grafts in the *recovered* group, both clinical motor scores and non-motor symptoms persisted following NP graft in the *non-recovered* group (**Figs.1B, S1**), in agreement with the absence of change in markers of striatal DA function and DAT (**Fig.3B**).

**Fig. 3.**
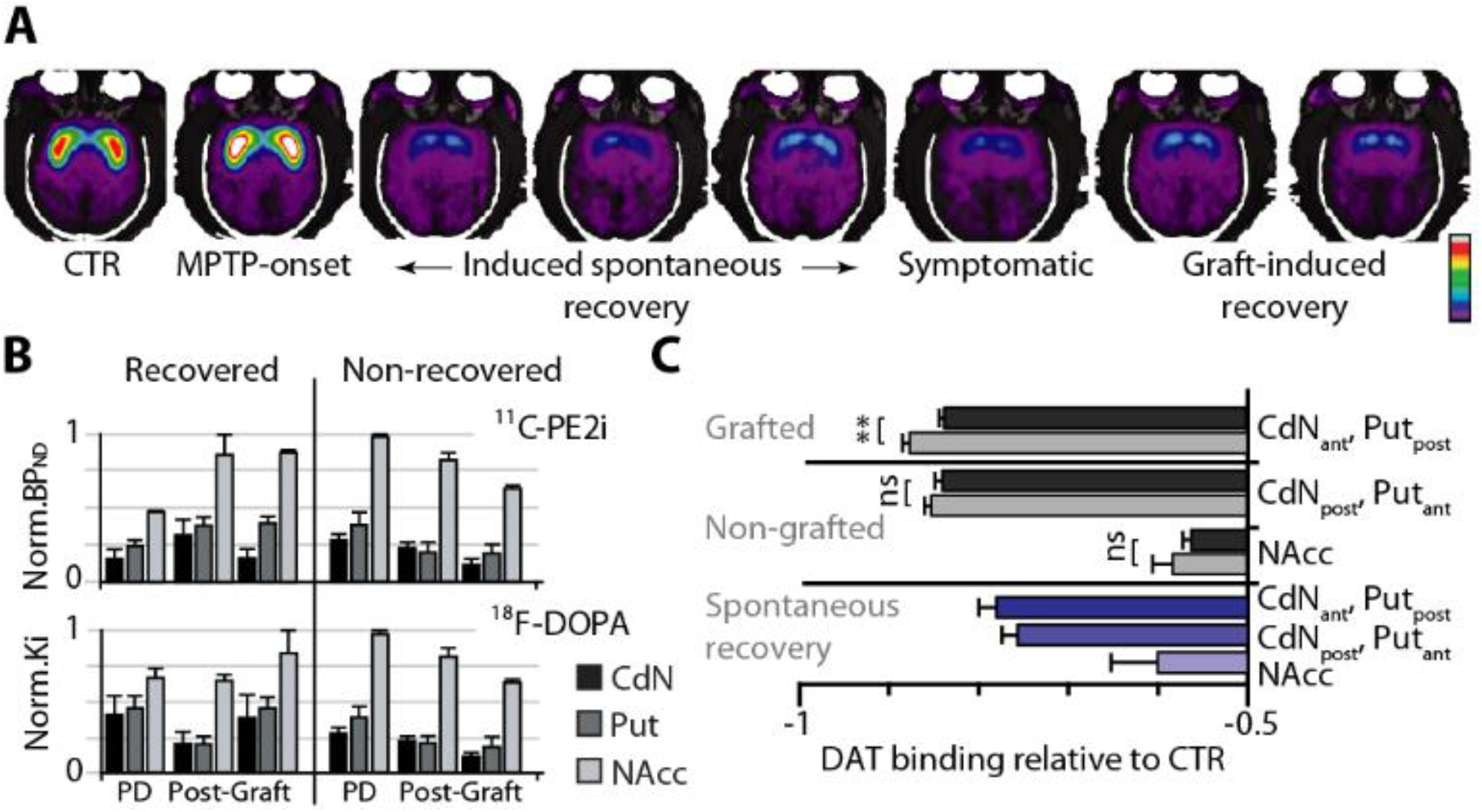
Functional markers of striatal DA function and DA terminals. (**A**) Images of striatal DAT binding potential (BP_ND_, color scale 0-7, case1 – see **Fig.S3A** for all cases). (**B**) Functional markers of nigrostriatal innervation (normalized BP_ND_ of ^11^C-PE2i) and DA function (normalized Ki, uptake of ^18^F-DOPA) in a *recovered* and a *non-recovered case*) during expression of stable symptoms (PD) and following transplantation (Post-Graft, minimum of 10weeks between measures). Min-max normalization per case. (**C**) Percent change in Striatal DAT binding in Grafted and Non-grafted regions for *recovered* (case1, black) and *non-recovered* (cases 3 and 6, grey). Purple, Striatal DAT binding following induced spontaneous recovery (case1). Two-sided Wilcoxon rank sum test, Bonferroni correction for multiple comparisons ** p < 0.01, Z = 2.92.

In the *recovered* group; we observed that PET-scan estimation of nigrostriatal innervation as revealed by DAT binding significantly increased bilaterally in the ventral striatum as early as 15 weeks post-graft **(Fig.1C**). This suggests a primarily protective influence of grafts on the remaining pool of nigrostriatal projection neurons. Post-graft F-DOPA scans confirmed a selective effect of NPs on DA synthesis (**Fig.3B**) that was associated with maximal motor recovery. Significant increases in DA function and DAT binding were also observed in grafted parts of the striatum when compared to the same regions in *non-recovered* cases (**Fig.3C**).

We performed sham transplantation (see **Materials and Methods**), in two out of the four cases following PET-scan imaging of striatal DAT innervation. No recovery from motor nor from non-motor symptoms was observed following sham grafts (**Fig.S3B**). Further, functional markers displayed no difference between *recovered* vs. *non-recovered* cases in striatal DAT binding and no specific increase in sham-grafted cases **(Fig.S3C**).

### Graft-induced recovery differs from spontaneous recovery

We compared the time course of behavioral symptoms and functional markers following graft-induced recovery to those following spontaneous recovery. In half of the cases (n=3), spontaneous recovery was initially induced by halting MPTP intoxication as described previously (Stage I, **Fig.1A**) [25, 27]. Spontaneous recovery of clinical motor symptoms displayed faster dynamics and was more complete compared to graft-induced recovery (large effect-size: *F*(2,13) = 24, *p* < 2.2*E*-16, 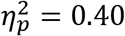, **Fig.4A**). Importantly, spontaneous recovery was not accompanied by changes in cognitive performance which contrasted with graft induced recovery that was invariably accompanied by progressive cognitive improvements (**Fig.4B**). These findings are intriguing because spontaneous motor recovery occurs despite an average 71.5±9.7% reduction in striatal DAT relative to control levels and that stable symptoms were observed when the lesion reached 82.3±7.9% of control [27]. The present findings show that graft-induced recovery occurs despite an average 78.04 ±12% reduction in striatal DAT binding (**Fig.3C**). These results confirm the specificity of graft-induced recovery compared to spontaneous recovery, particularly with respect to cognitive deficits (large effect-size for both Success: *F*(2,5) = 39, *p* < 2.2*E*-16, 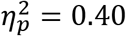 and Errors *F*(2,5) = 35, *p* < 2.2*E*-16, 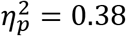, **Fig.4B**).

**Fig. 4.**
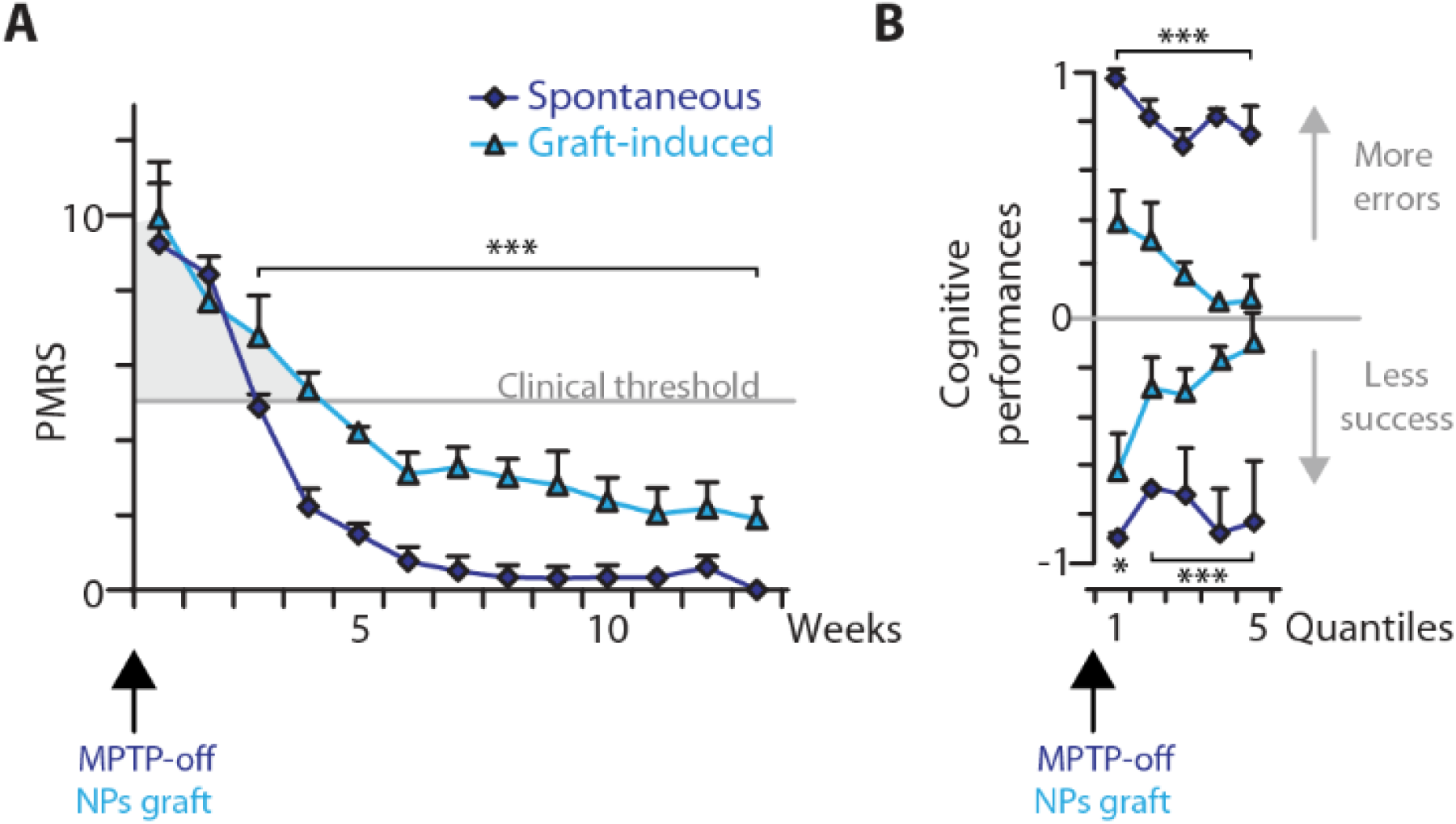
Dynamics of spontaneous vs. graft-induced behavioral recovery. (**A**) Clinical motor score over weeks and (**B**) cognitive performances (over quantiles) comparison between spontaneous recovery (cases 1, 3 and 4) vs. graft-induced recovery (cases 1, 4, 5). Mean ±SE, anova between groups, adjusted p-values for multiple comparison * < 0.05, *** < 0.001.

### Behavioral and functional recovery correlate with graft survival

We grafted NPs bilaterally in multiple sites including the ant-CdN, post-Put and SN. Each site received 1.5×10^5^-1×10^6^ NP cells (see **Materials and Methods**). Following an average post-graft period of 30±8weeks, we performed a *post-mortem* immuno-histological assessment of tau-GFP^+^ cells survival and integration into the host milieu (**Fig.1A**).

Noteworthy, the *non-recovered* group is characterized by a complete absence of tau-GFP^+^ cells (**Fig.S4**). This could be due to the fact that the nigrostriatal DA lesion was most severe in these cases (**Fig.2C**). However, immunological rejection of the grafted NPs cannot be excluded as no immunosuppressive treatment was employed. The fact that non-motor symptoms initially improved in the *non-recovered* group prior to an abrupt deterioration after the 2^nd^ Quantile (**Fig.1B**) is coherent with a chronic graft rejection in the *non-recovered* group (**Fig.S4**).

In sharp contrast, we found surviving grafted cells in all cases of the *recovered* group, at locations including the targeted functional domains of the basal ganglia (**Figs.5, S5-8**). The average number of surviving cells was: 1.23±0.9×10^4^ cells in the ant-CdN, 1.95±1.1×10^4^ cells in the post-Put and 5.93±3.0×10^4^ cells in the SN (**Table S3**) which corresponds to an average survival rate of: 6.9±6.2% in the ant-CdN, 13.0±7.3% in the post-Put and 15.5±9.4% in the SN (overall survival rate of 11.6±4.4%, Mean±SE).

**Fig. 5.**
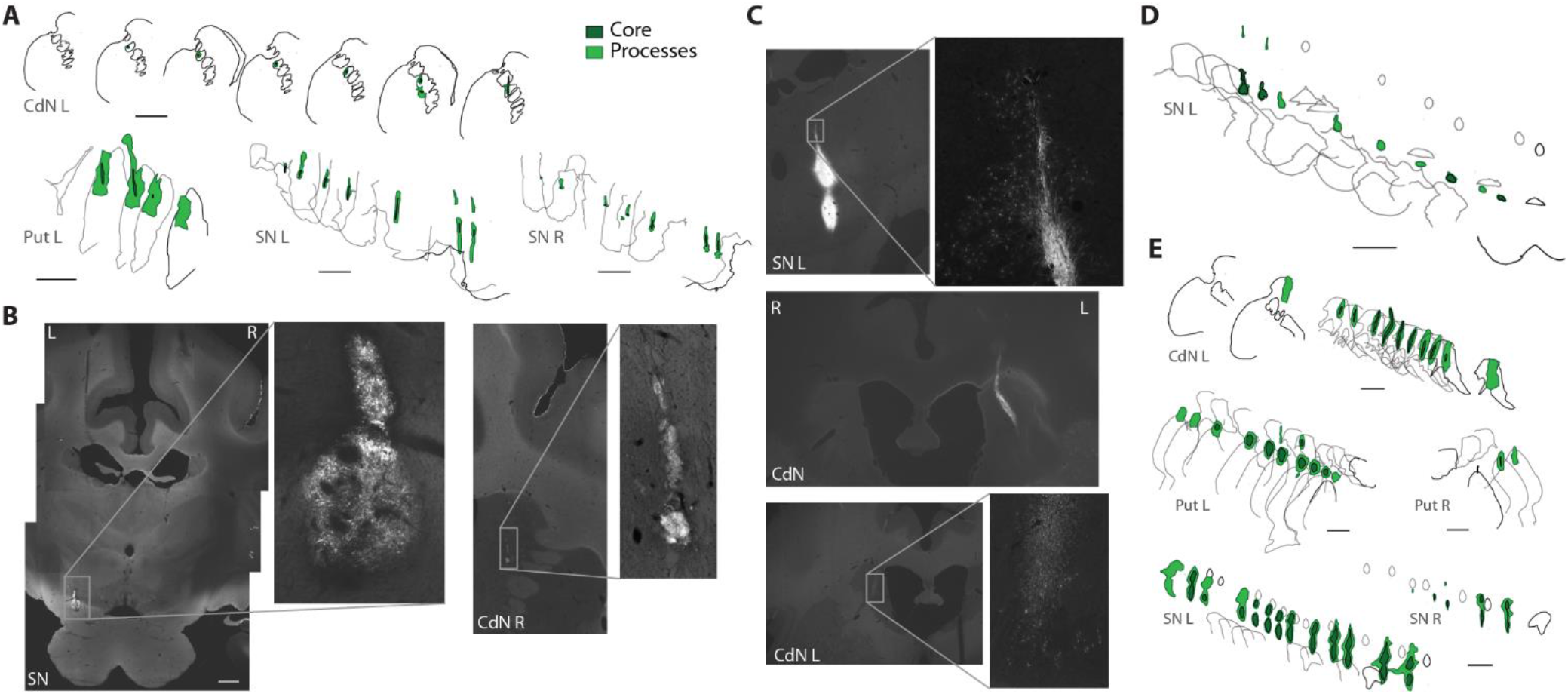
Post-mortem localization of grafted NPs. (**A**) Digital reconstructions of post mortem grafts in case1 (case4 shown in **D** and case5 in **E** - scale bars 200μm). (**B-C**) Photomicrographs of brain slices with zoom on tau-GFP^+^ grafts (scale bar 2mm, case4 shown in **B** and case5 in **C)**.

We observed unilateral survival of NPs in some of the grafted striatal regions (**Table S3**). We could not link asymmetrical survival in the post-Put to a systematic difference in left vs. right motor recovery (**Fig.S9A**) suggesting the induction of known cross-hemispheric compensatory processes [36]. Nevertheless, small differences were observed in cognitive performance linked to unilateral survival in the ant-CdN (**Fig.S9B**); differences that were coupled with a minor unilateral DAT binding decrease in the corresponding striatal region **(Fig.S9C**, right).

Surviving grafts displayed extensive tau-GFP positive projection fibers (**Fig.6A-C**) which extend several millimeters from the graft core, considerably further than the local processes which are less than 1 millimeter (**Fig.6D**). These findings indicate that the *recovered* group is characterized by integration of NPs into the host tissue (**Fig.6**).

**Fig. 6.**
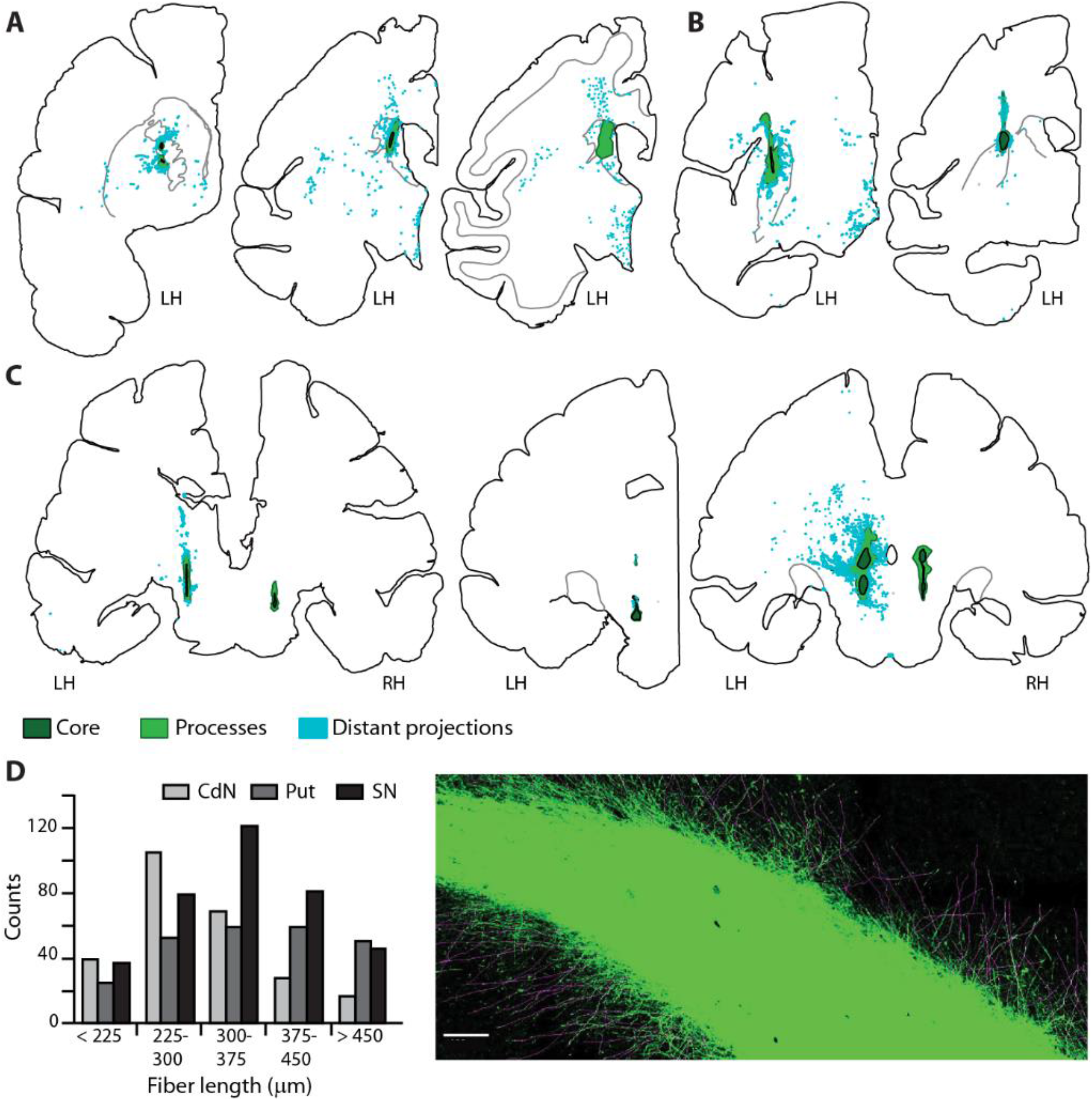
Grafted NPs project over long distances. Digital post-mortem reconstructions of NP grafts at the level of (**A**) anterior CdN in case1 (left) and case5 (middle and right), (**B**) posterior Put for cases 1 (left) and 5 (right) and, (**C**) SN for cases 1, 4 and 5 (from left to right). Tau-GFP+ projection fibers (clear blue) are found at relatively distant locations from the graft core (dark green), much farther than local graft processes (clear green) emanating from the core. (**D**) Typical processes emanating from the core have an average length superior to 340μm. Left, count histogram of processes’ length for case5. Right, typical tau-GFP+ photomicrograph showing graft-core and processes at the level of ant-CdN (case5, faint purple lines from measurement overlay, scale bar 100μm).

### Differentiation fates indicate protective and restorative effects of NP grafts

Detailed immuno-histological examination of graft locations reveals that grafted NPs differentiated principally into astrocytes and mature neurons (**Fig.7A-C, D; Fig.S5**). Among surviving NPs, astrocytic fate accounted for 42±5% (GFAP^+^/tau-GFP^+^ co-expression ratio, **Fig.7A**, **Table S4**) and neuronal fate for 16±4% (MAP2^+^/tau-GFP^+^ co-expression ratio, **Fig.7A**, **Table S4**). Furthermore, we demonstrate that NPs spontaneously differentiate into aminergic (TH+/DAT+) cells thereby potentially restoring a significant proportion of the lost DA pool (**Fig.7C-D**).

**Fig. 7.**
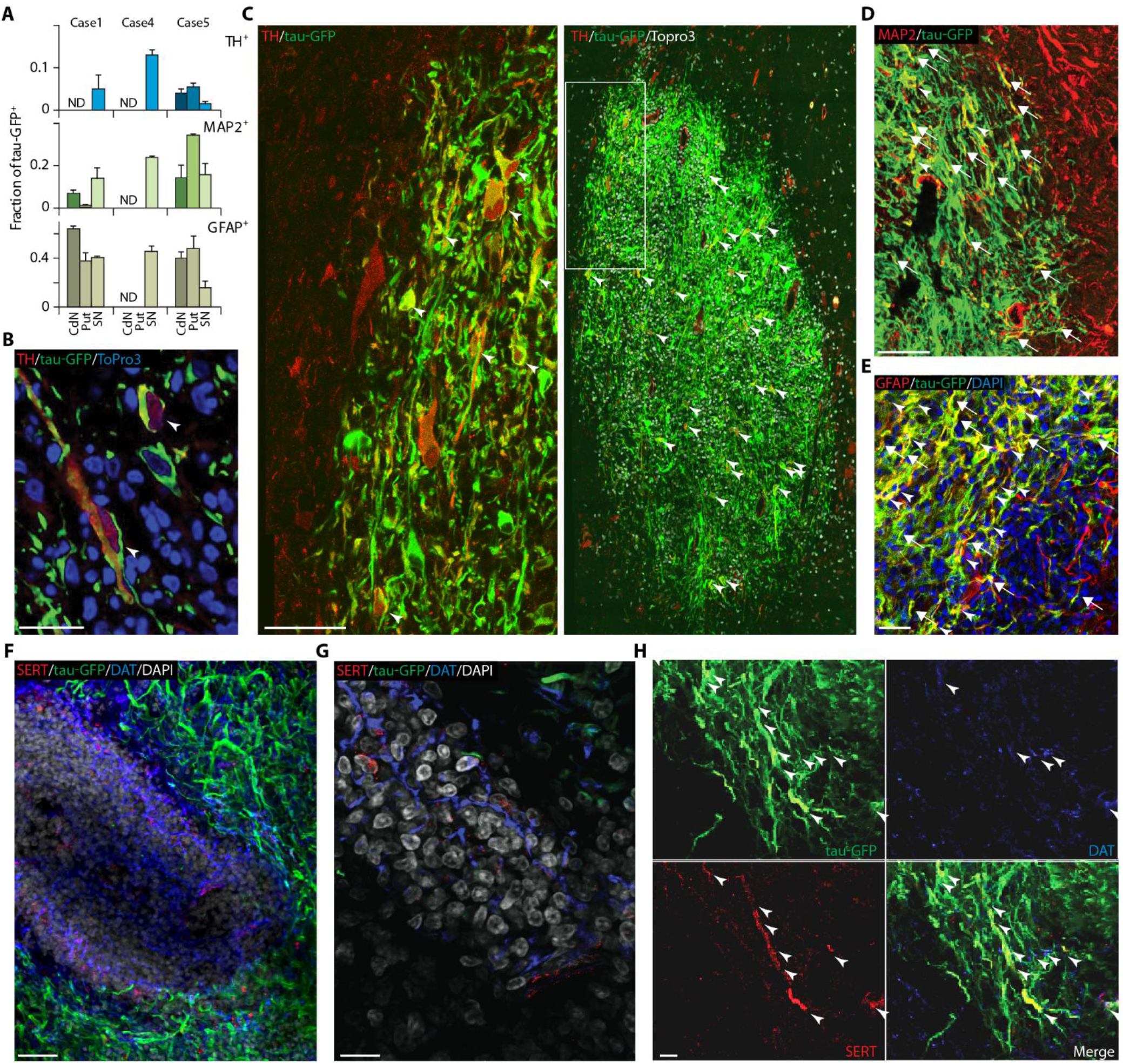
Differentiation Fate of grafted NPs. (**A**) Fraction of tau-GFP^+^ co-localized with TH^+^, MAP2^+^ and GFAP+. Illustrative example of NPs differentiated into: (**B)** aminergic cells, SN, case4 (arrowheads); (**C**) aminergic cells, SN, case5 (arrowheads, left is enlarged view of region framed on the right); (**D**) neuronal cells (arrowheads) and processes (arrows), post-Put, case5; (**E**) astroglial cells (arrowheads) and processes (arrows), SN, case4. (**F-G**) Host tissue within the graft core presenting numerous dopaminergic (DAT^+^, blue) and sparse serotoninergic (SERT^+^, red) processes. (**H**) NPs also display serotoninergic and/or dopaminergic phenotype (arrowheads). Scale bars, **B** 20μm, **C** left 50 μm a, **D-E** 100μm, **F** 25μm, **G** 5μm, **H** 15μm.

On average, the TH^+^/tau-GFP^+^ co-expression ratio implies that 6±2% of tau-GFP^+^ cells differentiated into aminergic cells (**Fig.7B**, **Table S4**). Up to 13% of tau-GFP^+^ in SN graft co-localize with TH^+^ expression (case4), which corresponds to an estimate of 1.4×10^4^ TH^+^/tau-GFP^+^ cells and hence to a restoration of nearly 18% of the original pool of DA-containing midbrain neurons in the same species [37]. We qualitatively confirmed NP differentiation into both the serotoninergic and the dopaminergic phenotypes (**Fig.7H**). Together with the protective effect of the graft on the survival or upregulation of DA cells (**Figs.1C, 7F-G**), these results strongly support a role of the graft in the recovery from motor and cognitive symptoms (**Figs.1**, **S1** and **Tables S4-S5**).

At some graft sites, aggregates of host cells (*i.e*. tau-GFP negative, **Figs.S7E, G and S8D, H**) were observed in the graft-core of nigral transplants, exhibiting aminergic TH^+^, dopaminergic DAT^+^ and sparse serotoninergic SERT^+^ processes (**Figs.7F-G, S7G-J**). Importantly, no post-graft overgrowth was observed (**Fig.S10A-C**) and few CD68^+^ macrophages were detected in the vicinity of transplanted sites, despite the absence of immunosuppressive treatment (**Fig.S10D-F**).

Finally, we explored the extent to which the grafted NPs were structurally integrated, via contacts with host neurons. For this purpose, we examined synaptophysin (SYN) immunohistology. SYN is localized in the membrane of pre-synaptic vesicles and provides a specific and sensitive marker for synapses. We explored regions at the periphery of the grafts, where the numerous local processes emanating from the core appear preferentially directed toward host DA cells (**Fig.8A, C**). Confocal microscopy shows that SYN co-localized at the junction between TH^+^ and MAP2^+^ host-cells and tau-GFP^+^ processes from grafted-cells (**Figs.8E, S8E, G, I-K**), indicating that local processes stemming from the graft core establish synaptic contacts with the remaining host DA cells in the SN (**Fig.8**). Together with the presence of long-distance projections emanating from the grafted sites (**Fig.6**), this suggests functional integration of NPs into the host brain circuitry and supports the effect of NP grafts in the observed functional recovery (**Figs.1B-C, 3**).

**Fig. 8.**
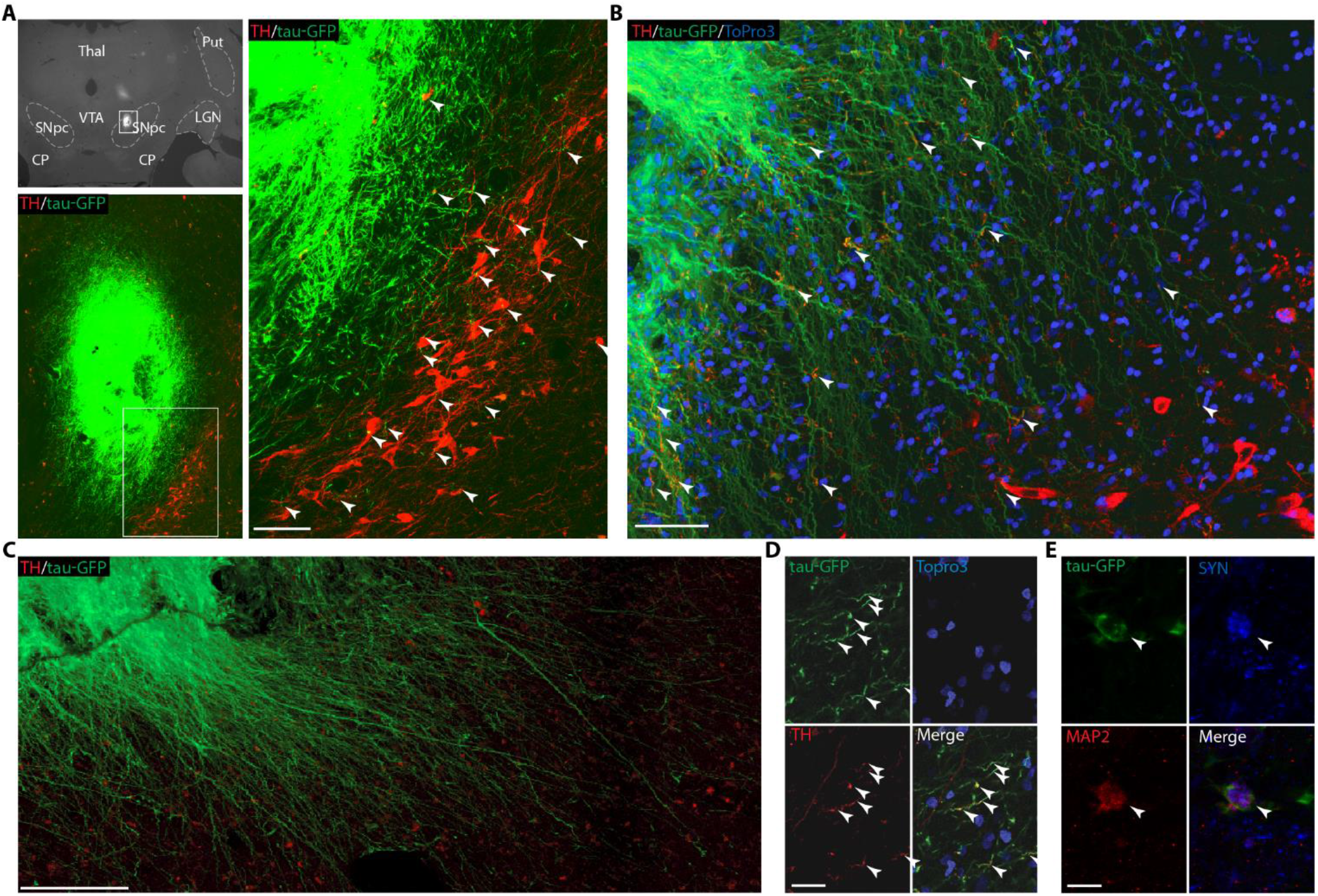
Histological evidence of NP integration and interaction with host cells. (**A**) Photomicrographs (up-left panel) shows position of graft in SN, framed region zoomed in bottom-left panel. Right-panel show zoom of framed region in the bottom-left panel. In (**A**) and (**B**), Grafted NPs (tau-GFP^+^, green) extend processes (arrowheads) toward surviving host DA cells (SN grafts). (**C**) NPs display numerous extensive processes projecting outside the graft-core. (**D**) NPs differentiate into aminergic phenotype displaying TH^+^ processes (arrowhead). (**E**) Synaptophysin expression (SYN, blue) co-localized with grafted cell (tau-GFP^+^) differentiated into mature neuron (MAP2^+^) suggesting synaptic contact. Scale bars: **A-C** 100μm, **D-E** 5μm.

## DISCUSSION

We transplanted macaque NPs into multiple functional domains of the basal ganglia, in both hemispheres of middle-aged parkinsonian macaques. Our study demonstrates that the successful integration of the grafts promotes a robust clinical and unprecedented cognitive recovery (**Figs.1B, S1**), as well as partial circadian recovery. Recovery is accompanied by quantitative effects on functional markers of nigrostriatal DA innervation and striatal DA function (**Figs.1C, 3B-C**). This study thus provides major empirical evidences in favor of a promising stem-cell derived cell therapy for PD patients. First, we show for the first time that transplantation and successful engraftment of NPs efficiently promotes recovery from motor and non-motor symptoms. Second, we demonstrate the restorative and the neuroprotective potential of grafted NPs. Our results show that NPs have the capacity to restore a significant pool of mature neurons (**Fig.7A**), including dopaminergic and serotoninergic cells. In addition, NPs largely differentiate into astroglial cells that the present study confirms provide a neuroprotective environment for the surviving pool of DA neurons [22–24]. Third, our results also suggest that nigral transplantation of NPs provide neurotrophic effects on the host cells (**Figs.7F-G, S7E, G-J and S8D, H**), orienting endogenous cells towards a DA fate [38, 39]. Finally, we provide cellular evidences of structural and functional integration of grafted NPs (**Figs.6, 8 and S8E, G, I-K**) into the host brain network.

The consistent cognitive recovery following successful engraftment of NPs in ant-CdN and SN establishes the potential of cell therapy to alleviate non-motor cognitive symptoms in addition to motor symptoms in PD patients. The specificity of the graft-induced cognitive recovery (**Fig.1B**) was further confirmed by the absence of cognitive improvements following sham-grafts (**Fig.S3B**) and spontaneous motor recovery (**Fig.4B**). These results anticipate the outcome of cell therapy for the majority of PD patients experiencing mild cognitive impairment (MCI) that significantly affects their quality of life [40]. Cognitive functions in human and NHP depend on the integrity of the prefrontal cortex (PFC), the DA system and the DA innervation of PFC via direct projections from SN and ventral tegmental area [41–46]. In addition, fronto-striatal circuits are also extensively involved in cognitive function and more particularly the cognitive sectors of the striatum i.e. ant-CdN [47]. Many cognitive symptoms expressed in PD patients not only arise from DA degeneration but also from other monoamine neurotransmitter systems closely connected to the DA system i.e. serotonin and norepinephrine [48, 49]. These systems are also affected in the MPTP-monkey model [50]. With the cognitive task that was used here, increased errors and reduced success rates observed in Stages I and II can be directly related to the reduced control over motor planning and parallel executive dysfunctions observed in PD patients [4, 51]. Hence, the observed reduction of cognitive performances (**Figs.1B, S1B**) can be directly imputed to the nigrostriatal DA lesion, the altered DA innervation of frontal cortex [41] and the wider MPTP-induced alteration of the aminergic system including noradrenalin and serotonin [29, 52–55]. We relate the consistent cognitive recovery following NP grafts (**Fig.1B**) to the efficiency of the transplanted NPs in both SN and ant-CdN of which an average of, respectively 5.93±3×10^4^ and 1.23±0.9×10^4^ cells survived, differentiated (**Fig.7**) and integrated the host network through local and long-distance projections (**Figs.6, 8**). Importantly, we show that grafted NPs can differentiate into a substantial proportion of TH^+^ expressing cells (**Fig.7A**), well within the range of previous studies reporting clinical improvement (**Table S4**), but also in the serotoninergic phenotype (**Fig.7H**) thereby supporting their role not only in motor but also in cognitive recovery.

In addition to these restorative effects, our results suggest that bilateral multi-site NP grafts exert a neuroprotective influence on the pool of surviving DA cells and of their ventro-striatal target. We show a significant and specific increase in striatal DAT binding following graft (**Fig.1C**) and not following sham-surgery (**Fig.S3C**). This increase is located in the ventral striatum that corresponds to the surviving pool of nigrostriatal innervation following induced spontaneous recovery [27]. The large proportion of astroglial differentiation (**Fig.7A**) might support this protective effect of NP grafts, in accordance with previous reports on human NP grafts in MPTP-monkeys [56]. Although we report some degree of recovery of DAT binding in the grafted caudate nucleus and putamen (**Fig.3B-C**), these effects were minimal compared to what was found in the ventral striatum. Longer post-graft survival could lead to greater increases in DAT binding. Indeed, transplanted PD patients display fast clinical improvements in motor function i.e. within 2-3 months after transplantation, whereas it can take 1-2 years to detect via functional imaging significant increase in DA function at the grafted locations [57–62].

The impact of cell therapy on chronobiological rhythms as far as we know has not been previously studied in the MPTP-monkey model. The initially observed circadian recovery following NP grafts concerned both groups but was followed very quickly by a deterioration after the third quantile in *non-recovered* cases. However, in the *recovered* group the LD ratio recovery was significant but only partial (**Fig.1B**). Previously we showed that following MPTP-induced nigrostriatal DA lesion, environmental cues mask the failure of the intact SCN to drive striatal clock genes and DA functions that control rest-wake locomotor rhythms [26]. Here, the partial recovery of circadian rhythms in the presence of environmental timing cues (**Fig.1B**), could reflect incomplete restoration of DA function. However, following NP grafts, under continuous light conditions (n=4, cases 1, 3, 4 and 5) there was an absence of endogenous circadian rhythm as observed in the MPTP monkey [26]. This again points to NP grafts failing to fully restore the DA network involved in the SCN control of rest-wake rhythms. DA projection neurons from SN and VTA regulate sleep-wake rhythms through interaction with several structures from the circadian network including the locus coeruleus, the lateral hypothalamus and the pedunculopontine nucleus [63]. Those structures, which were not grafted in the present study, are impacted in PD and following MPTP intoxication and thus constitute future candidates for cell replacement protocols [64–66].

There are limitations in the use of cell-therapy for PD e.g. presence of Lewy-body formations in the grafted cells [67, 68]. However, the balance benefit-risk of cell-therapy, even in cases of Lewy-body development in the grafted cells, is clearly in favor of the transplanted patients [11]. Three decades after the first clinical trials of transplanting cells from fetal ventral mesencephalon in PD patients [69], cell replacement therapy in PD is still not a proposed treatment. However, this therapeutic approach has recently been shown to be efficient at different levels [61, 62, 70], leading to more recent trials [71]. In parallel, many strategies have been developed in recent years in order to improve the safety and efficiency of using ESCs or iPSCs as a source for the production of the to-be-transplanted cells [21]. Virtually no NHP studies have addressed the impact of cell replacement on non-motor circadian symptoms and only one recent reported beneficial effects on depressive symptoms (**Table S5**). Here we report for the first-time efficient recovery from cognitive and clinical motor symptoms, as well as partial circadian recovery following NP grafts in the NHP model of PD. The present study confirms the neuroprotective and restorative capacities of NP grafts and, underlines the crucial importance of longitudinal and multifaceted approaches for a better translation to the clinic and appraisal of early-phase clinical trials.

While there are doubtless limitations in the use of the MPTP-monkey model to study human-specific aspect of PD; this model remains nevertheless the gold standard for modeling PD and for the pre-clinical evaluation of new therapeutic approaches. Previous studies have shown that advanced nigrostriatal lesion is detrimental to the survival of grafted cells [17, 18], which is coherent with our observation of poor survival of grafted NPs in the *non-recovered* group. Further, in the *non-recovered* group immune rejection is likely to have occurred as previously observed following allogenic transplantation in non-immunosuppressed monkeys [72, 73]. These observations suggest that future therapeutic cell replacement procedures should be carried out preferably at early stages of PD.

## MATERIALS AND METHODS

In agreement with the 3Rs [74], the rationale for the current study is that each case represents its own control through detailed follow-up of all the periods of the protocol i.e. CTR, MPTP, post-MPTP and Pre- vs. Post-Graft. We published precise and detailed reports for each clinical, behavioral and functional parameter followed as well as for the characterization of NPs grafted in the present study. These precise and detailed control procedures are described in **Table S1**.

### Study Approval

All procedures were carried out according to the 1986 European Community Council Directives (86/609/EEC) which was the official directive at the time of experiments, the French Commission for animal experimentation, the Department of Veterinary Services (DDSV Lyon, France). Authorization for the present study was delivered by the “Préfet de la Région Rhône Alpes” and the “Directeur départemental de la protection des populations” under Permit Number: #A690290402. All procedures were designed with reference to the recommendations of the Weatherall report, “The use of non-human primates in research”.

### Study Design

The objective of the research was to investigate the efficiency of NP grafts to restore motor and non-motor symptoms in parkinsonian macaque monkeys. All behavioral and functional measures were acquired by experimenters blinded to the outcome of the grafts integration as this was evaluated post-mortem. Post-mortem quantifications of graft survival, differentiation rate and length of graft’s processes were done by investigators blinded to the precise evolution of parkinsonian symptoms of the animals following graft. The experimental findings include replication of the procedure in all individual cases separately and conclusions on the findings arise from difference in outcome based on post-mortem evaluation of grafts integration. Cases followed by PET imaging were chosen randomly before experiment starts as well as cases chosen for sham-grafts. Data collection was stopped at least 7 months following transplantation and after last possible PET-scan acquisition. Data-points of the locomotor activity acquired on the day of anesthesia for MRI, PET-scan or surgery were excluded from circadian analyses. Presented results include all other data-points, no outliers were excluded.

Nine macaque monkeys (Macaca fascicularis, n=9) were used in total for this study. The study was designed such that one third (n=3) were randomly selected as histological controls i.e. never received MPTP injections and, the other two third (n=6) entered the MPTP protocol. Whilst we selected control animals for histological comparison, we designed the study such that each animal would be its own control for clinical, behavioral and functional measures. **Figure 1A** displays a schematic representation of the study design and protocol. Two third of the MPTP group (n=4) were randomly selected to be followed by PET imaging of striatal DA transporters and DA function throughout the full study and, half of them (n=2) were randomly selected to evaluate sham-grafts effects on clinical, behavioral and functional measures. In addition, half of the animals entering the MPTP protocol were randomly selected (n=3) for a two-step MPTP procedure in which MPTP injections were first halted based on clinical score, in order to induce spontaneous recovery [25] for further comparison of this *spontaneous* recovery to the prospective *graft-induced* recovery. MPTP-intoxication was then resumed in order to induce stable motor symptoms [27] over the period preceding cell therapy. Study design is further detailed below.

Six late middle-aged – 11-13 (13-17) years old at protocol onset (end) female macaque monkeys (Macaca fascicularis, 4-5kg) were intoxicated with low-dose 6-methyl-1-Methyl-4-phenyl-1,2,3,6-tetrahydropyridin injections (MPTP, 0.2mg/kg, i.m). Animals were housed in a room dedicated to MPTP experiments, with free access to water and received food twice a day. The neurotoxin was delivered at low-doses chronically each 3-4 days (slowly progressive lesion) during prolonged periods, followed by acute low-doses intoxication (daily injections) in order to ensure stable persistent motor symptoms, as described previously [27, 29]. Cases 1, 3 and 4, were selected to critically compare the evolution of clinical, behavioral and functional markers following spontaneous clinical motor recovery [25, 27] with those obtained during potential recovery induced by cell therapy. In order to do so, we suspended the slowly progressive lesion induced by chronic low-dose MPTP injections as soon as the clinical score reached symptomatic threshold (**Fig.S1** and **Table S2**). In all cases, acute (daily) low-dose MPTP injections induced persistent motor-symptoms. We cautiously stopped daily MPTP injections when the PMRS-motor score was above ten for two consecutive days following one MPTP injection. For results presentation, animals were grouped according to their clinical motor state at the end of the protocol following cell therapy (see **Fig.1**) i.e. 1^st^ group – recovered (n=3, cases 1, 4, 5) and, 2^nd^ group – non-recovered (n=2, cases 3 and 6). Case2 had per operative brain hemorrhage following NPs transplantation and was thus excluded from group comparison. Data are presented according to the following periods of the protocol: 1-CTR (measures acquired before MPTP-onset); 2-MPTP (during MPTP intoxication period); 3-post-MPTP (following arrest of last MPTP injections, clinical motor score remained stably above 5 during this period i.e. motor-symptomatic); 4-post-Grafts (following transplantation, described in **Supplementary Materials and Methods**). Delays between last MPTP injection and cell therapy were 4-23 weeks (n=6, all cases) and, delays between last MPTP injection and sham-grafts were 6-16 weeks (n=2, cases 1 and 6).

Complete details concerning Parkinsonian Motor Rating Scale – PMRS, Surgical Procedures, Transplanted cells, Cognitive behavior – detour task, Circadian rhythm follow-up, Dopamine function imaging – [^11^C]-PE2I and [^18^F]-FDOPA, Immuno-histological and Quantification procedures, Statistics can be found in the **Supplementary Materials and Methods**.

## Acknowledgments

We acknowledge Dr. Vincent Leviel for discussions at different stages of the study. We thank Frank Lavenne for PET acquisition; Didier Le Bars and Frédéric Bonnefoi for radioligand synthesis; Mélissa Remy, Léo Pion and Luc Grinand for histological graft analyses; Claude Gronfier for advice during circadian analyses; Pascale Giroud for precious help during anesthesia, surgery and perfusion; and Kenneth Knoblauch for help in statistical analyses.

## Funding

The funders had no role in study design, data collection and analysis, decision to publish, or preparation of the manuscript.

Région Rhône-Alpes (C.D, J.V)

Fondation de France (E.P, H.M.C, J.V)

Fondation Caisse d’Epargne Rhône-Alpes Lyon (C.D, J.V)

Rhône-Alpes cible 11-010869 (H.K)

Fondation Neurodis (C.R.E.W)

Cluster Handicap Vieillissement Neurosciences Rhône-Alpes and LabEX CORTEX : Agence Nationale de la Recherche ANR-11 LABX-0042 (H.K, E.P), P6-2005 IST-1583 (H.K), FP7-2007 ICT-216593 (H.K)

Agence Nationale de la Recherche ANR-11-BSV4-501 (H.K).

## Author contributions

Conceptualization: HMC, EP, HK, PS, CD, JV

Methodology: HMC, EP, HK, PS, CD, JV

Investigation: FW, KD, KF, CREW, AB, VD, PM, CL, JV

Visualization: FW, JV

Funding acquisition: CREW, HMC, EP, HK, CD, JV

Supervision: CD, JV

Writing – original draft: FW, HK, JV

Writing – review & editing: FW, KF, CREW, HMC, EP, HK, PS, CD, JV

## Additional Information

Correspondence and requests for materials should be addressed to JV.

## Supplementary Materials

Supplementary Material and Methods.

Fig.S1. Individual follow-up of behavioral and clinical markers.

Fig.S2. Characterization of grafted cells.

Fig.S3. Functional follow-up of striatal DAT binding and Sham-grafts consequences on behavioral and functional markers.

Fig.S4. Grafted NPs did not survive in the *non-recovered* group.

Fig.S5. Post-mortem localization and characterization of transplanted NPs.

Fig.S6. Grafted NPs in case1.

Fig.S7. Grafted NPs in case4.

Fig.S8. Grafted NPs in case5.

Fig.S9. Hemispheric differences in graft-induced recovery.

Fig.S10. NPs show no overgrowth after transplantation and induce minor immune reaction.

Table S1. Longitudinal protocol design, repeated measures and case controls.

Table S2. Total cumulated MPTP doses per case.

Table S3. Cell survival following transplantation of NPs.

Table S4. Grafted cells number, survival rates, differentiation fates and comparison to literature.

Table S5. Clinical motor, cognitive and functional impact of grafts described in Table S4.

Table S6. Antibodies list.

Table S7. Primers for semi-quantitative RT-PCR.

Table S8. Group-level stats of Fig.1B.

