## Supplementary Materials for "Induced Cognitive Impairments Reversed by Grafts of Neural Precursors: a Longitudinal Study in a Macaque Model of Parkinson’s Disease"

### SUPPLEMENTARY MATERIALS AND METHODS

#### Parkinsonian Motor Rating Scale – PMRS

Clinical scoring was done independently by three trained experimenters at separate times throughout the day, 3-5 days per week using the Parkinsonian Monkey Rating Scale [25-27]. The PMRS contains a motor component, employing a previously published scale [75] used to define the onset of the motor symptomatic period. The total score for freezing (0–3), resting tremor (0–3, for right and left side), rigidity (0–3), posture (0–2), bradykinesia (0–3), and ability to manipulate food (0–3, for right and left side) was used to frame injections (maximum total score of 23): a score below 5 defined the non-symptomatic state in premotor and recovery periods, and a score above 5 defined symptomatic periods. During the symptomatic post-MPTP period, all animals displayed the following clinical motor symptoms: freezing of gait, resting tremor, stooped posture, bradykinesia and difficulty to manipulate food at varying degrees. Only two cases displayed rigidity (cases 1 and 6), that was confirmed by clinical examination i.e. arm manipulation. No further measure than observation was done to evaluate resting tremor and it might have been indistinguishable from postural tremor at rest [50]; however, resting tremor is a consistent feature in NHP after chronic low-dose MTP intoxication [25, 29, 76, 77] while acute MPTP intoxication induces tremor only in some NHP species [78, 79]. Case 4 had a high-score post-MPTP (motor symptoms>15), characterized by tremor at rest and freezing. This motor-symptomatic state persisted during the 4 weeks prior to transplantation (**Fig.S1A**). Case 6 had a high-score post-MPTP (motor symptoms>15), characterized by severe rigidity. Musculoskeletal pain is a recognized source of pain syndromes and discomfort in Parkinson's disease patients. We thus delivered regularly ketoprofen (Profenid), from week5 post-MPTP, in order to relieve suspected apparent musculoskeletal pain. Subsequently the clinical score for this animal progressively improved, due to amelioration on the rigidity scale and ability to manipulate food (from week5 to week10 post-MPTP, **Fig.S1A**), but stabilized at week10 and remained largely symptomatic (score above 12) despite continuous treatment and disappearance of apparent musculoskeletal pain syndrome. Case 6 was transplanted at week23 post-MPTP.

#### Surgical procedures

We performed all transplantations bilaterally using 5µL Hamilton syringe and small trepanations over target sites. Stereotaxic coordinates were calculated from individuals T1 and T2 structural MRI images. We undertook sham-grafts (trepanation and injection of 5µL PBS without NPs) in cases 1 and 6 in order to control for behavioral and clinical effects due to the surgical act in itself. We placed sham-grafts (PBS-only) in posterior caudate and anterior putamen, using the same protocol as for NPs transplantation. The maximum number of injected cells was $1\times10^6$ and, the minimum number of injected cells varied between $3\times10^5$ for SN sites and $1.5\times10^5$ for striatum sites (**Table S4**). For all transplantations, the syringe was lowered 500µm further than the target position and then retracted to target; cells were then injected at a rate of 1µL/min after a 5min delay. Following a 3min delay after injection completion, the syringe was retracted slowly by 2mm and, another 2min delay was observed before full retraction. We assessed cell viability by trypan blue exclusion before and after injection by counting the remaining cells. Viability at both time points exceeded 95%.

#### Transplanted cells

We derived neural precursors (NPs) from a rhesus embryonic stem cell (RhESC) line stably expressing tau-GFP (LYON-ESC line) [31]. NPs were obtained either as described in [80], and amplified in the presence of EGF and FGF2 (NPs, **Fig.S2A**, upper panels) or after MS5 induced-neural differentiation, as described in [81] followed by early midbrain DA differentiation (mDA-NPs, **Fig.S2A**, lower panels). RT-PCR analysis returned similar marker expression profiles (except for the specific DA markers LMX1A and LMX1B which are not expressed in NPs; **Fig.S2A-B**). Detailed protocols are further described below. Therefore, no a priori distinction between NPs derived from the two protocols was made for transplantation in different brain locations.

To isolate NPs, RhESCs were manually selected and co-cultured on mitomycine C (MMC)-treated mouse embryonic fibroblasts ($2.5\times10^4$cells/cm$^2$) in KO-DMEM medium containing Knockout serum replacement (KO-SR), non-essential amino acids (NEAA, 1X), and β-Mercaptoethanol (0.1mM), for 7-13 days as described in [80]. Neuroepithelial-like cells that spontaneously emerged were selected manually (Passage 1 – P1) and cultured in DMEM/F12 medium containing N2 supplements (1X), non-essential amino acids (NEAA, 1X), β -Mercaptoethanol (0.1mM), Fibroblast Growth Factor2 (FGF2, 20ng/mL, Merck Millipore) and Epidermal Growth Factor (EGF, 20ng/mL, Sigma Aldrich). Amplification was done through mild trypsinization (Trypsin 0.025% - EDTA) followed by soybean trypsin inhibitor treatment. Medium was changed every other day. To induce early DA midbrain differentiation, RhESCs were seeded on MMC treated MS5 cells (day0). NPs emerging after 11-13 days were manually selected, dissociated with trypsin-EDTA, and cultured on matrigel-coated dishes (BD Biosciences, San Diego, http://www.bdbiosciences.com) in Neurobasal medium containing B27 supplement (without Vitamine A, 1X), 5 µg/mL bovine fibronectin, FGF2 (20ng/mL). From day6-8, we added the dorso-ventralizing factors: FGF8 (100ng/mL), Sonic hedgehog (SHH, 200ng/ml) and FGF20 (2ng/mL) as described previously [81]. To induce differentiation into mature DA neurons, Wnt3a (100ng/mL) was added to the culture medium from day11 to day24. From day24 to day45, mDA-NPs were cultured in the presence of Brain Derived Neurotrophic Factor (BDNF, 20ng/mL, Sigma Aldrich), Glial cell line-Derived Neurotrophic Factor (GDNF, 20ng/mL, R&D Systems), Transforming Growth Factor β2 (TGFβ2, 2ng/ml, R&D Systems), dibutyryl cAMP (0.5mM, Sigma Aldrich), Neurotrophin-3 (NT3, 20ng/mL, Sigma Aldrich), Wnt5a (100ng/mL) and Ascorbic Acid (AA, 0.2mM, Sigma Aldrich). To induce neuronal differentiation, NPs were trypsinized (Trypsin 0.025% - EDTA, 0.1g/L), replated ($2.5\times10^4$cells/cm$^2$) into Matrigel-coated dishes, and cultured in DMEM/F12 medium supplemented with N2 (1X) and FGF2 (10ng/mL). After 7 days, the medium was changed to DMEM/F12 medium mixed with Neurobasal medium (1:1) supplemented with N2 (0.5X), B27 (0.5X) and FGF2 (5ng/mL). After an additional 7 days in these conditions, medium was switched to Neurobasal medium supplemented with B27 (1X). To induce glial differentiation, NPs were cultured in DMEM/F12 medium supplemented with 10% Fetal Bovine Serum (FBS, HyClone, Perbio) for 7 days. All reagents were from Invitrogen, unless otherwise indicated.

For cell transplantation, we dissociated NPs into single cells using trypsin-EDTA to prepare a suspension at a concentration of $0.3\text{-}2.0\times10^5$ cells/µL and, injected 5µL per site. Grafted NPs express specific neural markers (**Fig.S2A-B**) and showed *in-vitro* potential to fully differentiate along glial and neuronal pathways (**Fig.S2C**), including mature DA neurons (**Fig.S2D-E**). Despite the fact that RhESC-derived NPs were transplanted in Macaca fascicularis, we choose to not carry out immunosuppressive treatment as previously done in other Xenografts studies [82, 83] and, in preclinical studies assessing the potential of human cell grafts. Relying on the extensive sharing of MHC alleles [84] and hybridization evidence [85], we anticipated low immune response and decent survival [86]. We avoided the inconvenient of immunosuppressive therapy in Parkinsonian macaques (e.g. increased risk of infectious complications) by assuming limited immunogenicity of transplanted cells differentiated from RhESC into the cynomolgus macaque brain.

**Cognitive behavior – detour task**

Cognitive performance was monitored 3-5 days a week using a previously described behavioral test [25, 27]. Briefly, performance on the 'detour task' was evaluated by the percent of successes (retrieval of reward on the first reach, over the total number of trials) and errors (barrier hits, over the total number of responses observed - there could be several responses per trial except in the case of success). Errors due to motor impairments were classified as such and further discarded from the performance evaluation, so as to avoid confounds due to the symptomatic state of the animal. The performance on the 'detour task' depends on the integrity of frontal cortex, the dopaminergic system, the dopaminergic innervation of frontal cortex [87] and has been used to assess the impact of lesion of the nigrostriatal axis on cognitive performance as well as potential therapy for cognitive impairments [88-90]. For inter-individual comparison, percent errors and success were, for each case, first converted in percent change by the average control value and normalized between 0 and 1 by overall maximum and minimum score respectively.

**Circadian rhythm follow-up**

Animals were maintained under an LD cycle (12h light: 12h dark) of approximately 450–500 lux during the light phase and 0 lux during the dark phase. Locomotor activity was continuously recorded in 1 min intervals throughout the entire duration of the study using passive-infrared motion detectors mounted above each animal's cage. The motion captors were connected to a computerized data acquisition system (Circadian Activity Monitoring, INSERM, France [91]). Data were analyzed using the Clocklab software package (Actimetrics, Evanston, IL, USA). We previously used Chi-squared periodogram method (**Table S1**) to calculate the period of behavioral rhythm which is defined as the time elapsed for one complete oscillation or cycle (the distance in time between two consecutive peaks or troughs of a recurring rhythm). The stability and the consistency of the rhythm are reflected by the value of the periodogram amplitude which corresponds to the zenith of a periodogram curve. Non-parametric circadian rhythm analyses (NPCRA) were used to estimate the strength and fragmentation of rest-activity rhythms [25, 26]. These included measurements of inter-daily stability (calculated as the ratio between the variance of the average 24-hour pattern around the mean and the overall variance and gives an indication of the consistency of day to day activity or the strength of coupling to the LD cycle), intra-daily variability (calculated as the ratio of the mean squares of the difference between successive hours and the mean squares around the grand mean and gives an indication of the frequency of transitions between rest and activity periods, corresponding to the fragmentation of the rhythm) and, relative amplitude (ratio between acrophase and nadir of the rhythm, representing the ratio between activity amplitude in light and dark phases). When the rest-activity rhythm is stable, inter-daily stability is high, the intra-daily variability is low and relative amplitude is high (CTR, **Fig.S1**).

**Dopamine function imaging – [$^{11}$C]-PE2I and [$^{18}$F]-FDOPA**

To evaluate the in-vivo DA function, we acquired images from Positron Emission Tomography (PET) scans using (E)-N-(3-iodoprop-2-enyl)-2beta-carbomethoxy-3beta-(4'-methylphenyl)-nortropane labeled with carbon 11 ([$^{11}$C]-PE2I) at different intervals throughout the protocol as described previously [27] in cases 1, 2, 3 and 6 and in cases 1, 3 and 6 after grafts at different delays (see **Figs.S1, S3A**). [$^{11}$C]-PE2I specifically binds with high affinity and selectivity to DA transporters (DAT, Ki = 17nM) and is considered to provide an index of the integrity of the DA pathway that has been used for PD diagnosis, see [92-94] for a review. Additionally, we used images from PET scans using L-3,4-dihydroxy-6-($^{18}$F)fluorophenylalanine ([$^{18}$F]-FDOPA, an analog of the DA precursor L-DOPA) to evaluate the central dopaminergic function of pre-synaptic neurons, i) just before grafts (during stable expression of typical motor parkinsonian symptoms i.e. stage II, the symptomatic period) and, ii) after grafts at different delays (**Fig.S1**). We performed [$^{11}$C]-PE2I and [$^{18}$F]-FDOPA PET scans with an ECAT Exact HR+ tomograph (Siemens CTI), in 3D acquisition mode, covering an axial distance of 15.2cm. The transaxial resolution of the reconstructed images was about 4.1mm full-width and half maximum in the center. We acquired transmission scans with three rotating 68Ge sources to correct emission scans for the attenuation of 511 keV photonrays through tissue and head support. Full procedure for PET-scans acquisition, modeling and ROIs definition for regional assessment of [$^{11}$C]-PE2I changes have been published [27]. Fluoro-DOPA was labeled with $^{18}$F-fluoride (cyclotron–produced isotope, half-life = 109 min). Specific radioactivity was 3.27 ± 1.3 mCi/µmol. Radiochemical and chemical purity of produced [$^{18}$F]-FDOPA (as determined by HPLC) was above 99%. After anesthesia was induced and head secured with a MRIcompatible stereotaxic frame (Kopf, CA, USA) to reduce variability in the measure, a cannula was inserted in the femoral vein. [$^{18}$F]-FDOPA was injected as a bolus over a 4 s period followed by a saline flush. Radioactivity was measured in a series of 26 sequential time frames of increasing duration (from 30sec to 5min; total time 90min). PET modeling and regional assignment of PET changes were done as previously described [27] and applied to F-DOPA measurement [52]. Statistical positive (shown in **Fig.1C**) and negative (shown in **Fig.S9C**, right) difference were determined from 2.5$^{th}$ and 97.5$^{th}$ percentile boundaries of voxel-level distribution statistics of all brain ROIs as described previously [27] and illustrated in **Fig.S9C**, left.

**Immuno-histological procedures**

Deep anesthesia was induced with Ketamine after premedication with chlorpromazine hydrochloride (Largactil) followed by a lethal dose of pentobarbital sodium (Vibrac, 100 mg.kg-1, i.p.; confirmed by complete loss of corneal reflex) before animals were transcardially perfused with saline (0.9% with procaine), 4% paraformaldehyde and 0.05%-glutaraldehyde in phosphate buffer. Cryoprotection was ensured by sucrose gradients (10-30%) perfusion post-fixation. Brains were removed, kept in cryoprotecting liquid overnight and coronal 50µm thick sections cut on a freezing microtome and collected serially. Sections were processed together with equivalent sections from control animals for Tyrosine Hydroxylase (TH; Millipore, #MAB318) and DAT (Millipore, #MAB369) immunochemistry visualized with 3,3-diaminobenzidine, in $Ni^{2}$+H202 (0.5-1.0%). All sections were washed, mounted, dried, and dehydrated in increasing gradients of ethanol, cleared in toluene, mounted with mounting medium and cover slipped. Immunostainings of cultured cells were done according to previously published protocol [31]. Briefly, cells were fixed with 2% PFA in PBS at 4°C for 1 hour, and permeabilized in Tris buffered saline (TBS) with 0.1% Triton X-100 (3×10min). Non-specific binding was blocked with 10% normal goat serum (NGS) for 20min at room temperature (RT). Cells were incubated overnight at 4°C with primary antibodies diluted in Dako diluent. After three rinses in TBS, cells were exposed to secondary antibodies for 1 hour at RT followed by nuclear staining with DAPI (1/10,000) for 3min. After three rinses in TBS, coverslips were mounted on slides, and observed under confocal microscopy. For *post-mortem* immunofluorescent staining of brain slices, sections were washed six to ten times in Tris buffered saline (TBS) and permeabilized in Triton X-100 (0.5%, SIGMA, #T9284). Nonspecific binding was blocked with 10% NGS or donkey serum (Goat: Invitrogen; #16210064; Donkey: SIGMA #D9663) for 45 min at room temperature. Sections were incubated for three days at 4°C, with primary antibodies diluted in Dako antibody diluent (Dako #S3022) supplemented with 0.5% Triton X-100. They were then exposed to secondary antibodies and DAPI (1/10,000; Invitrogen #D1306) for 2 hours at RT. Sections were mounted on slides and examined using a confocal microscope (Leica TCS SP5). Sections were regularly processed for GFP immunostaining to detect grafted sites extent (every five sections i.e. every 250µm). Immunostaining of cultured cells were done according to previously published protocols [31] and the list of antibodies used presented in **Table S6**. For semi-quantitative RT-PCR analyses, total RNA was prepared with a QIAGEN RNeasy kit (QIAGEN, Valencia, CA, http://www.qiagen.com). Standard reverse-transcription reactions were performed with 1µg of total RNA primed with random primers using SuperScript II first strand synthesis system, according to the manufacturer's recommendations (Invitrogen). Oligonucleotide sequences are given in **Table S7**.

**Quantification procedures**

We measured the length of processes emanating from the graft core by measuring the length of tau-GFP positive projections (ImageJ, NeuronJ plugins, http://imagej.nih.gov/ij/), presented in **Fig.6D**. We estimated each graft volume through a 3D reconstruction process using contours (ExploraNova, Mercator, http://www.exploranova.com/products/mercator/), presented in **Figs.5, 6A-C**. We determined the average cell density by i) counting nuclei stained with ToPro3 or DAPI of tau-GFP$^{+}$ cells in four random locations per confocal plane i.e. confocal optical section, in several slices at different levels of the graft core to return cell density per $mm^3$ and ii) by scaling those counts to the estimated graft volume in $mm^3$ to return estimated cell number per graft. Finally, we calculated survival rate (**Table S3**) according to the number of transplanted cells per site (**Table S4**). To estimate the proportion of $TH^+$, $MAP2^+$ and $GFAP^+$ areas in each grafted site, confocal images were acquired at random locations of the tau-$GFP^+$ grafts. We measured $TH^+$, $MAP2^+$, $GFAP^+$ and tau-$GFP^+$ areas on binary images of the corresponding channel (ImageJ, http://imagej.nih.gov/ij/) and determined the co-expression ratios of $TH^+$/tau-$GFP^+$ $MAP2^+$/tau-$GFP^+$ and $GFAP^+$/tau-$GFP^+$ on at least 4 z-sections per grafted site. For semi-quantitative immunohistochemistry, three age and weight matched Macaca fascicularis served as controls. We evaluated the extent and amplitude of the nigrostriatal lesion, excluding graft locations as previously described [27, 95]. The mean optical density (O.D.) of TH and DAT immunopositive regions were computed on at least 5 sections per animal (5-20) from 8bit images, normalized and compared with controls (immunostaining with 3,3'-diaminobenzidine – DAB revealed in the same bath [27]) with two-sample t-test separately for each ROI (ttest2, Matlab). In **Fig.2B**, composite CTR and MPTP images has been enhanced for brightness and contrast, for illustration purpose. Importantly, decreased levels of striatal DAT O.D. compared to controls were consistent with the decrease of DAT as evaluated *in-vivo* by PET.

**Statistics**

We segmented data for MPTP, post-MPTP and post-Graft periods into five equal epochs and grouped variables into each of these epochs. This method of segmentation, described in [25, 27], thus reveals normalized stages of the progression of processes and allows for comparison of different parameters across an equivalent number of epochs for all subjects, called Quantiles (**Figs.1, 4B, S9A-B**, 27 ± 3 days for the CTR period; duration per quantile for premotor 10 ± 5 days , motor/pre-graft 18 ± 8 days and post-graft 42 ± 12 days). Results are presented as means ± standard errors. If not stated otherwise, parameters were compared between groups by a parametric one-way anova test of the difference between groups (estimated standard errors and t-values were computed using linear fit and group-contrast result was given by two-tailed p-values based on a t-statistic). We also report the effect-size of the anova test (range from 0 to 1, partial eta-squared, $\eta_p^2$). Effect-size is considered small for values < 0.09 and large for values > 0.25, moderate in between. Detailed statistics corresponding to significance tests in **Fig.1B** are reported in **Table S8**. Significance was considered at $p<0.05$. Correction for multiple comparisons was applied to adjust p-values using the Bonferroni method (rstatix package, R). Statistical analyses were computed using R software (R Foundation for Statistical Computing, Vienna, Austrian http://www.R-project.org) and the lm, anova and summary functions (Chambers, J. M. and Hastie, T. J. (1992) Statistical Models in S. Wadsworth & Brooks/Cole).

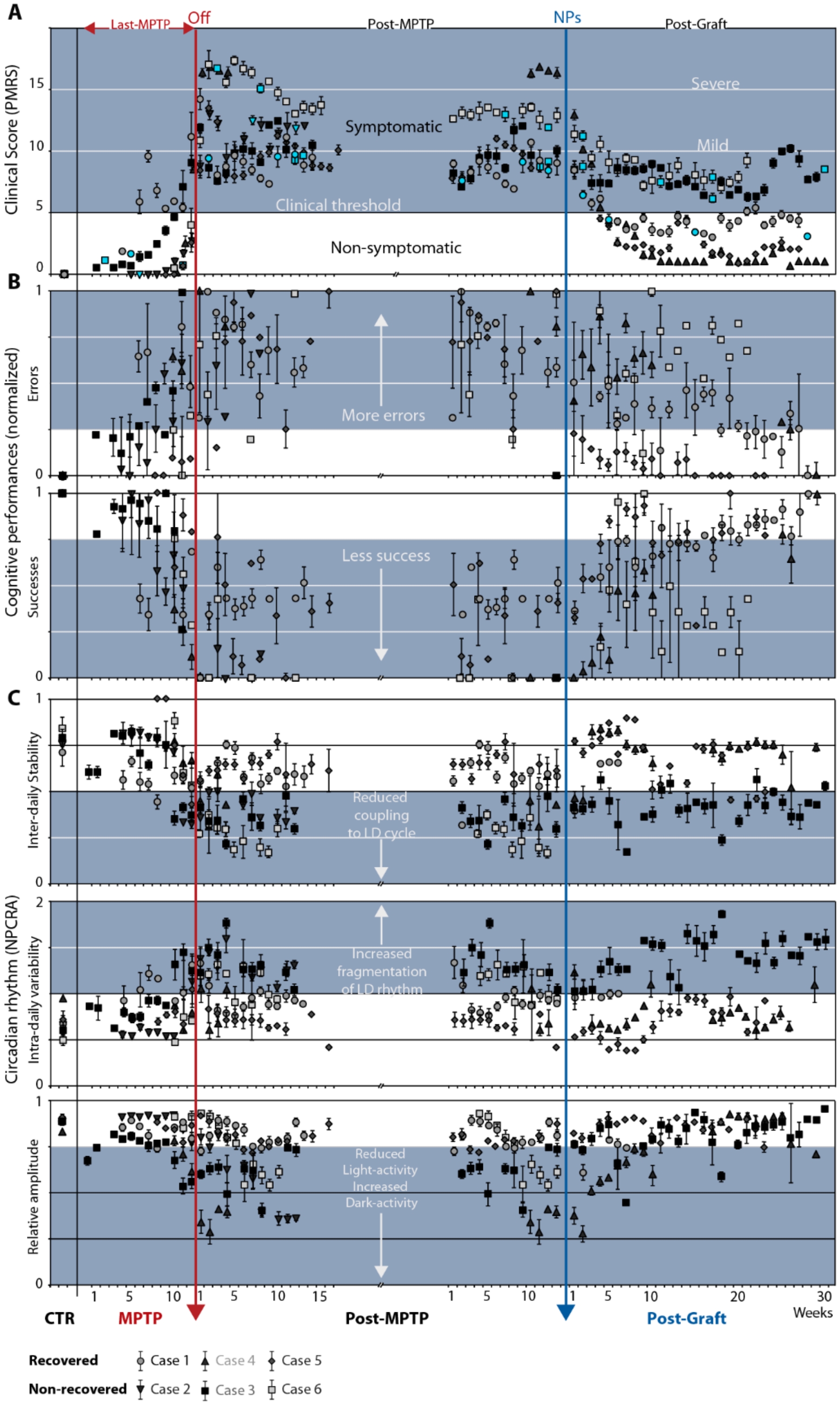


**Fig.S1: Individual follow-up of behavioral and clinical markers**

Motor and non-motor behavioral markers per case (legend down left) after alignment to MPTP offset (red line) and NP grafts (blue line). (**A**) Clinical motor score, a score ≥ 5 (clinical threshold) indicates symptomatic state corresponding to clinical diagnosis for PD patients. (**B**) Cognitive performances (Errors – up and Successes – down at the object retrieval detour task, see Methods for details), normalized by min-max per case. (**C**) Non-parametric Circadian Rhythm Analysis (NPCRA). Mean±SE. Markers colored in blue in panel (**A**) when PET-scan measures were acquired (day of measure excluded from weekly average).

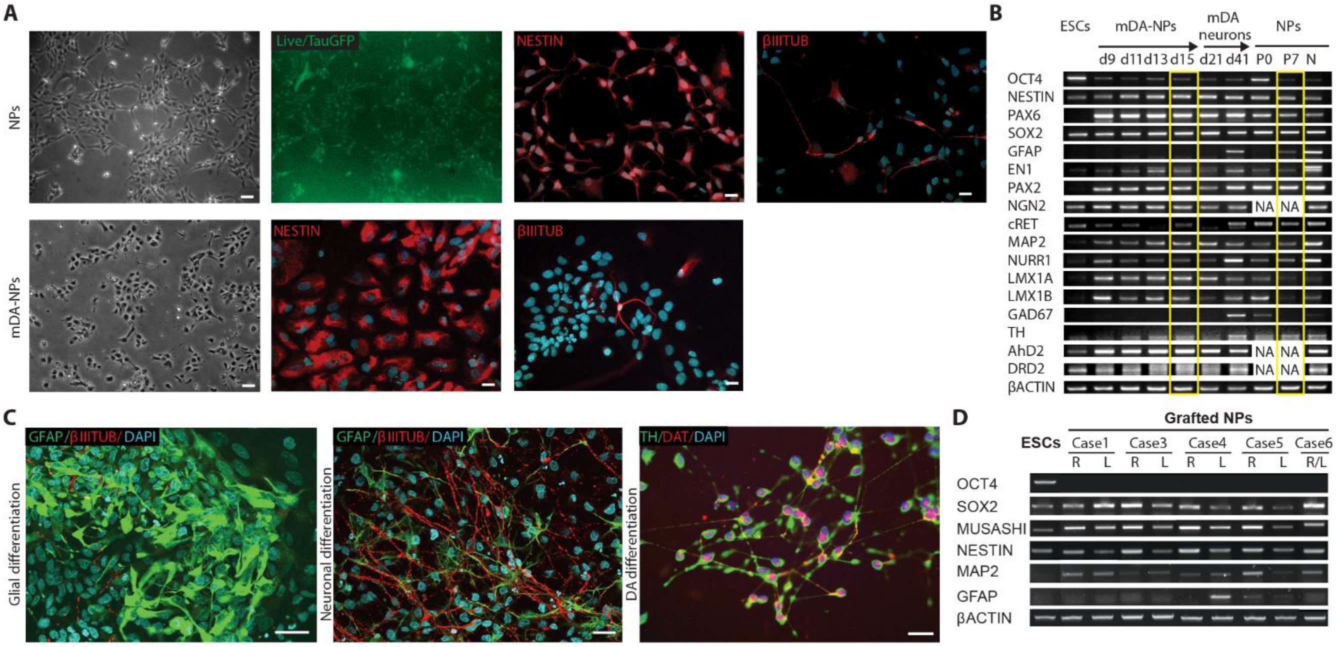


#### Fig.S2: Characterization of grafted cells

(**A**) NPs derived from the rhesus Embryonic Stem Cells (RhESCs, LYON-ES1 line) stably expressing tau-GFP. Upper panels from left to right, NPs at Passage 4 (P4), phase contrast and corresponding live fluorescence images (scale bar 50µm) and immunostainings for NESTIN and βIII-TUBULIN (scale bars 20µm). Lower panels from left to right, mDA-NPs isolated from rosettes (P1d17), phase contrast image and immunostainings for NESTIN and βIII-TUBULIN (scale bars 20µm). **(B)** From left to right, RT-PCR analysis of specific gene markers in ESCs, during differentiation into mDA-NPs and mDA-neurons, and in NPs (P0-P7 and N for NP-derived neurons). For example, markers specific for ESCs (OCT4, SOX2), NPs (NESTIN, SOX2, PAX6), glial (GFAP), neuronal (MAP2), DA progenitors (LMX1A, LMX1B, NURR1), or DA neurons (TH, AhD2, DRD2). Yellow rectangles indicate stages at which NPs were transplanted. (**C**) Immunofluorescence staining for βIII-TUBULIN and GFAP upon glial (left) and neuronal (middle) differentiation of NPs (scale bars 50µm) and, staining for DAT and TH of mDA-neurons (d40) (scale bar 10µm). (**D**) RT-PCR analysis showing comparison of marker genes in ESCs and NPs, for each case, at the time they were transplanted in right (R) and left (L) hemisphere. d, day; βIIITUB; βIII-TUBULIN.

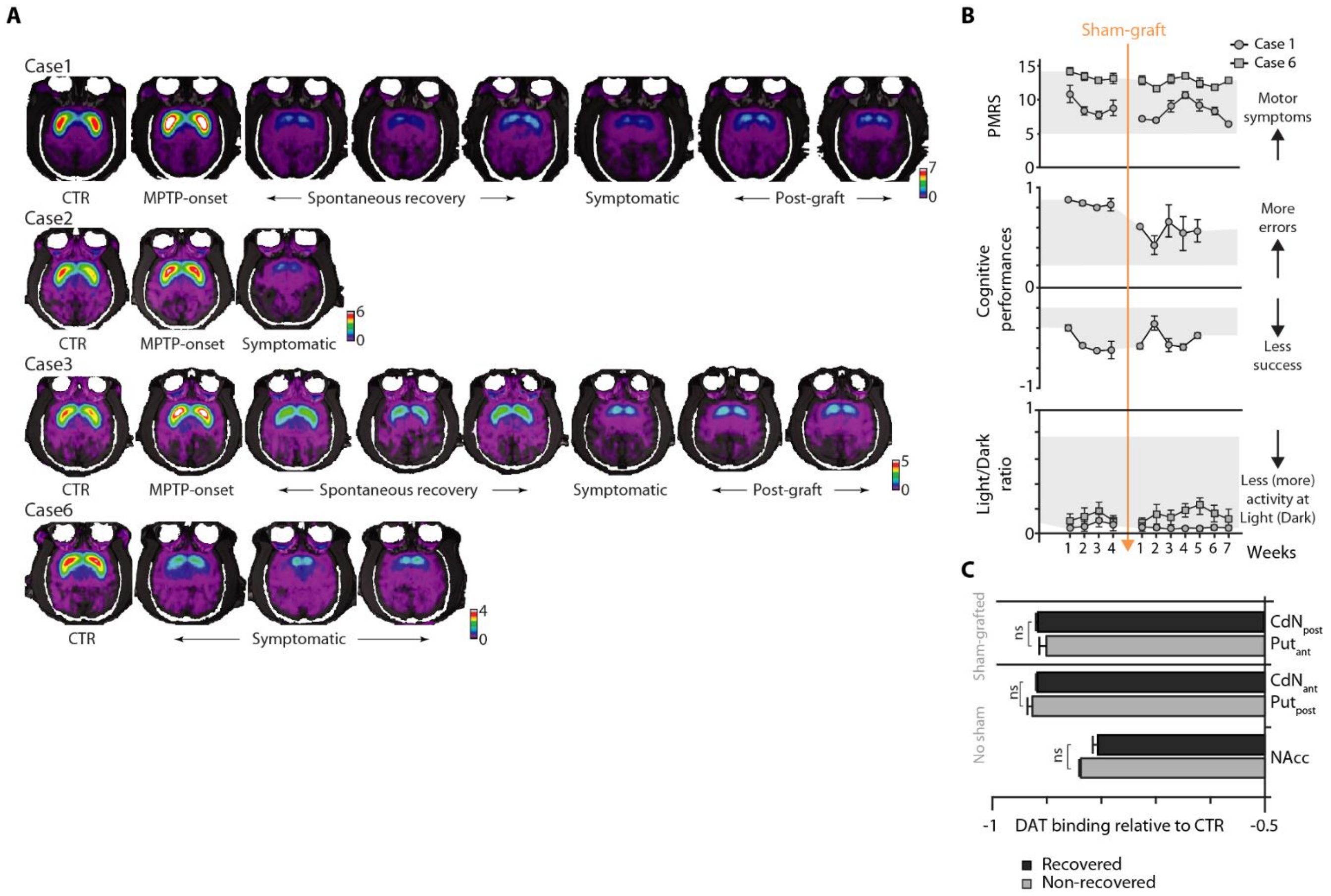


**Fig.S3: Functional follow-up of striatal DAT binding and Sham-grafts consequences on behavioral and functional markers.**

(**A**) PET-scan images reconstruction after alignment to individual MRI showing progression of DAT binding from CTR until post-Graft periods. Color scale for $BP_{ND}$. (**B**) Sham-grafts, performed in cases 1 and 6, show no effect on behavioral improvement. From top to bottom, clinical motor score – PMRS, cognitive performances (errors – up, successes – down, range normalized to control performances, -1 for success) and Light/Dark ratio. Means ± SE. (**C**) PET-scans of $^{11}C$-PE2I (relative change compared to CTR) show no difference in DAT binding following Sham-graft between *recovered* (case1) vs. *non-recovered* (case 6).

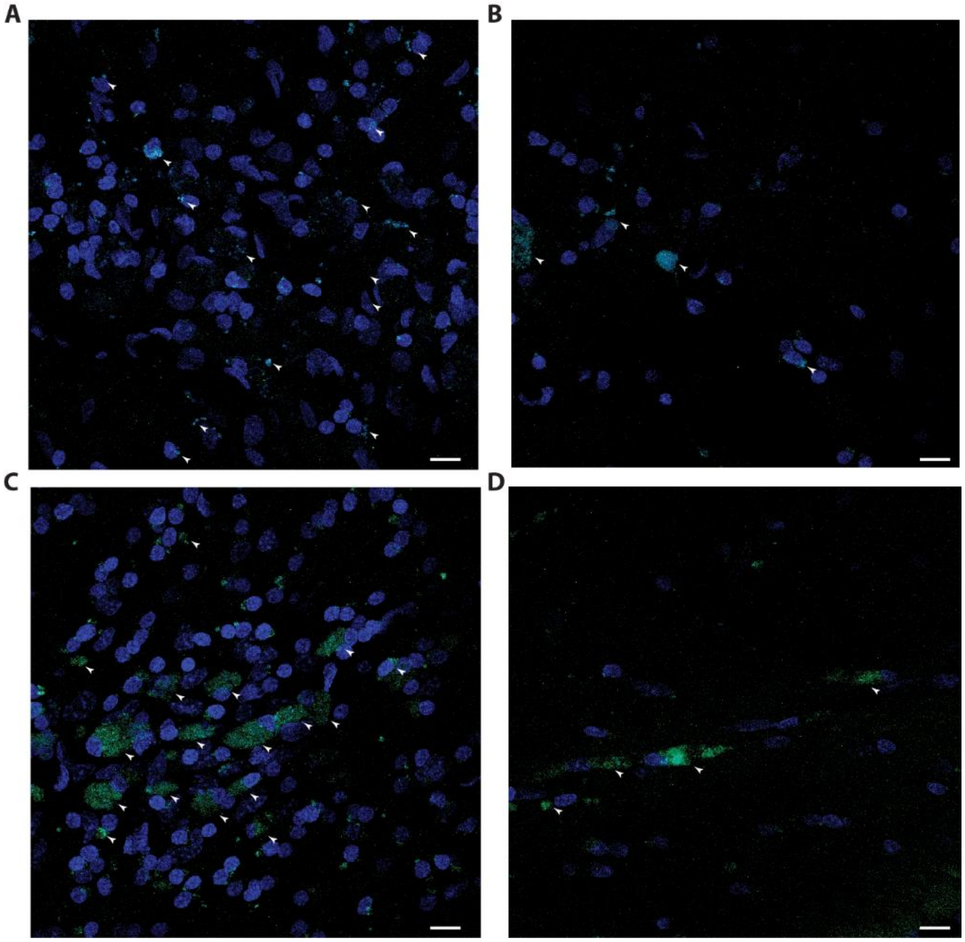


**Fig.S4: Grafted NPs did not survive in the *non-recovered* group.**

(**A-D**) tau-GFP (green) and ToPro3 (blue) immunolabeling of SN tissue at grafted locations in case 6 showing only cell debris (arrowheads) surrounded by weak and diffuse tau-GFP staining. Scale bars, 10µm.

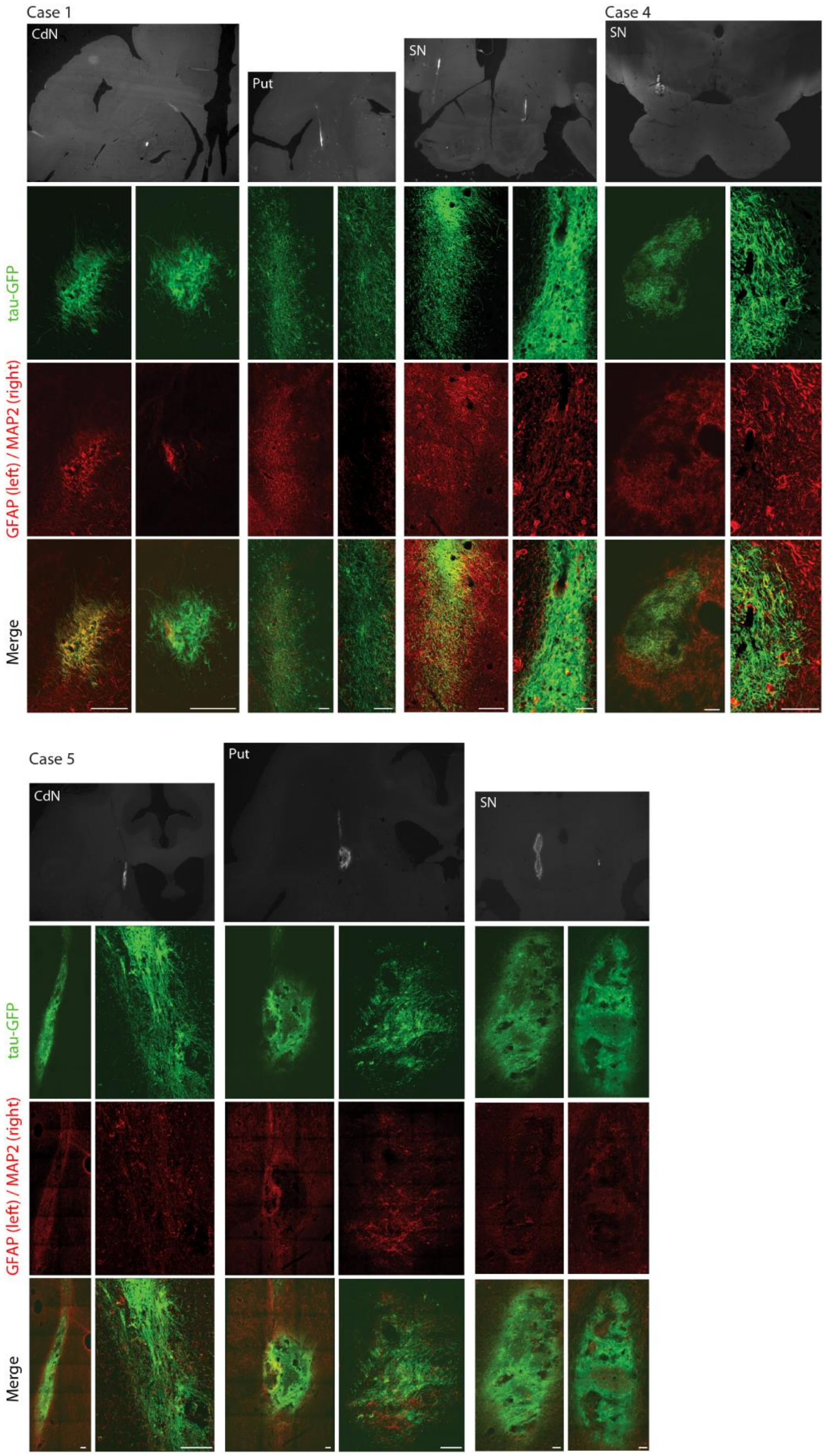


**Fig.S5: Post-mortem localization and characterization of transplanted NPs.**

Photomicrographs of tau-GFP$^{+}$ grafts and immunofluorescence labeling at different sites for cases 1, 4 and 5. Scale bars, 100µm. . Grayscale images from fluorescent microscope (filter 488nm, LeicaM165 FC).

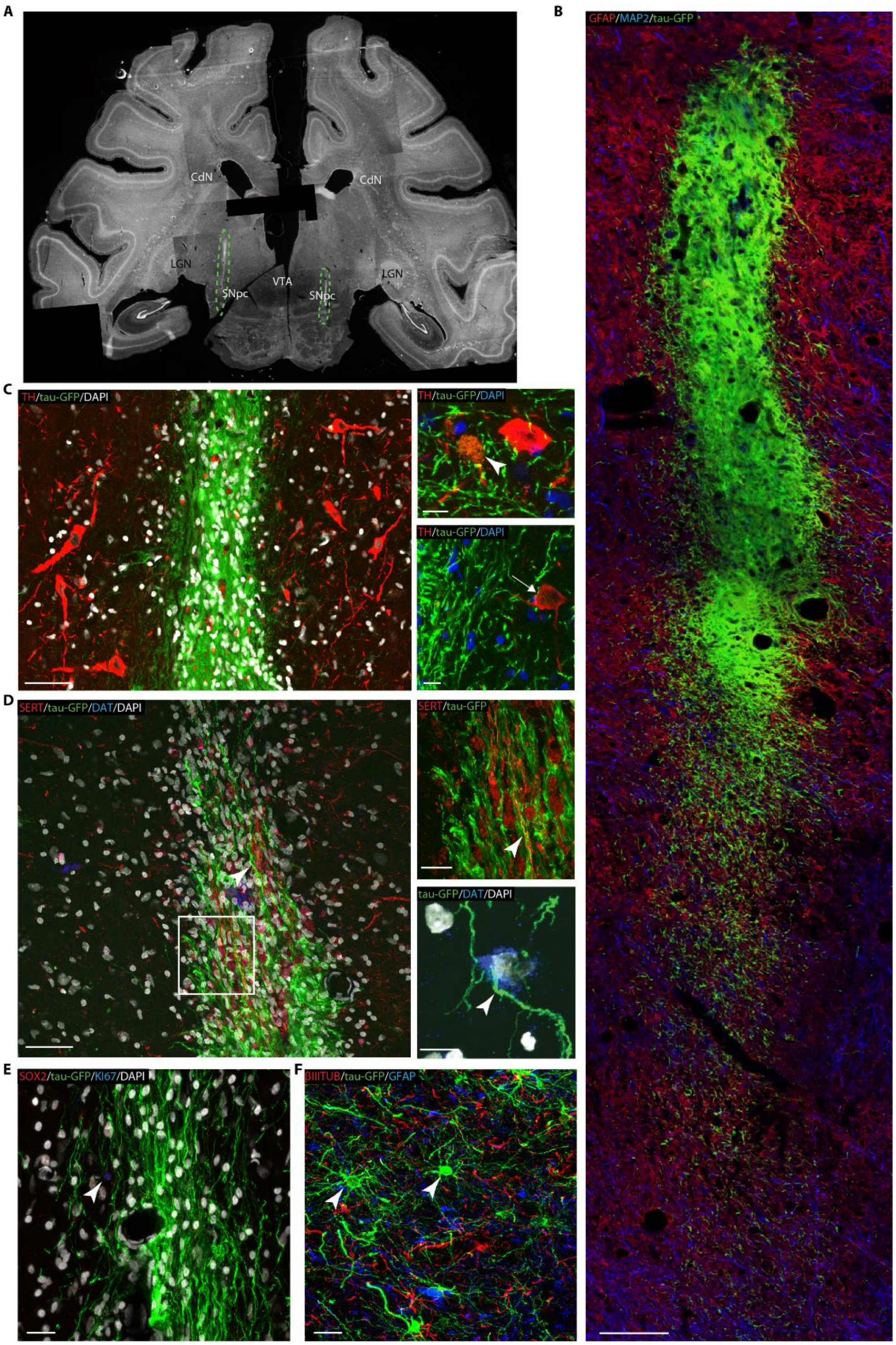


**Fig.S6: Grafted NPs in case 1.**

(**A**) Photomicrograph from fluorescent microscope acquisitions (tau-GFP$^+$ in white, filter 488nm, LeicaM165 FC) showing location of NPs in nigral transplant (green). (**B**) Closer view, confocal acquisition at an equivalent section. (**C**) Grafted cells (tau-GFP$^+$) in SN tissue surrounded by surviving host TH$^+$ cells, some of which co-express TH$^+$ (arrowhead, top-right panel) and seems to contact host aminergic cells (arrow, bottom-right panel). (**D**) Photomicrograft at SN site showing grafted cells processes co-expressing SERT$^+$ (frame enlarged in top-right panel, arrowhead) and surrounding host DA cell (arrowhead, DAT$^+$, bottom-right panel). (**E**) Host cell expressing KI67 (arrowhead). (**F**) Grafted cells showing glial morphology. Scale bars: B 100µm; C 10µm; D left 50µm, top-right 10µm, bottom-right 20µm; E-F 20µm.

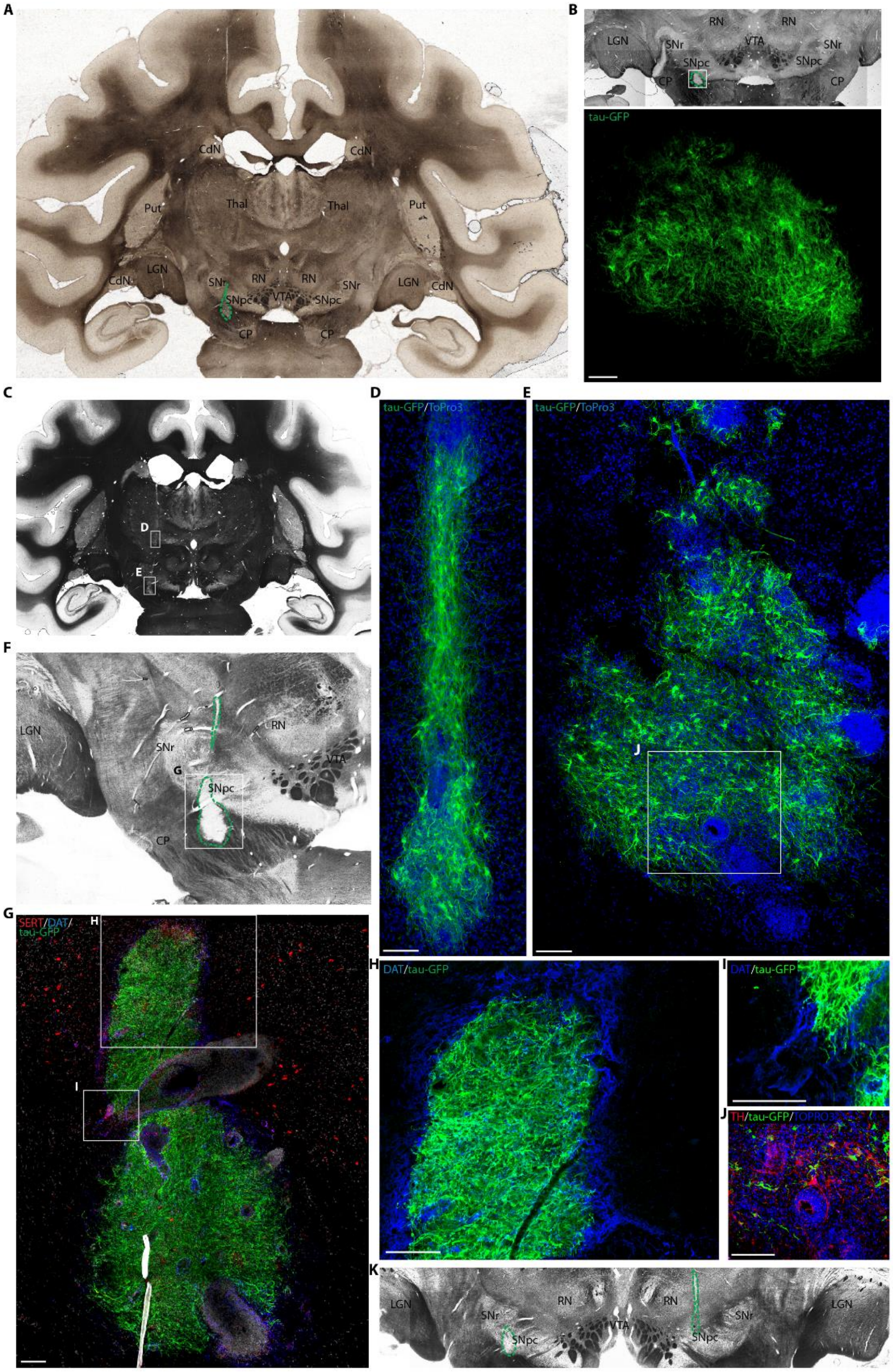


**Fig.S7: Grafted NPs in case 4.**

(**A**), (**B**), (**C**), (**F**), (**K**) Grayscale images from white-field microscope showing location of NPs in nigral transplant (green). (**B**) Bottom panel show tau-GFP$^+$ cells framed in top panel. (**C**) Framed regions shown in (**D**) and (**E**). (**G**) Framed region in (**F**) showing concentration of DAT+ processes on the graft core and surroundings, shown at higher magnification in (**H**) and (**I**). (**J**) Host cells aggregate surrounded by a host TH$^+$ cell, region framed in (**E**). Scale bars: B, E, G-J 100μm; D 50μm.

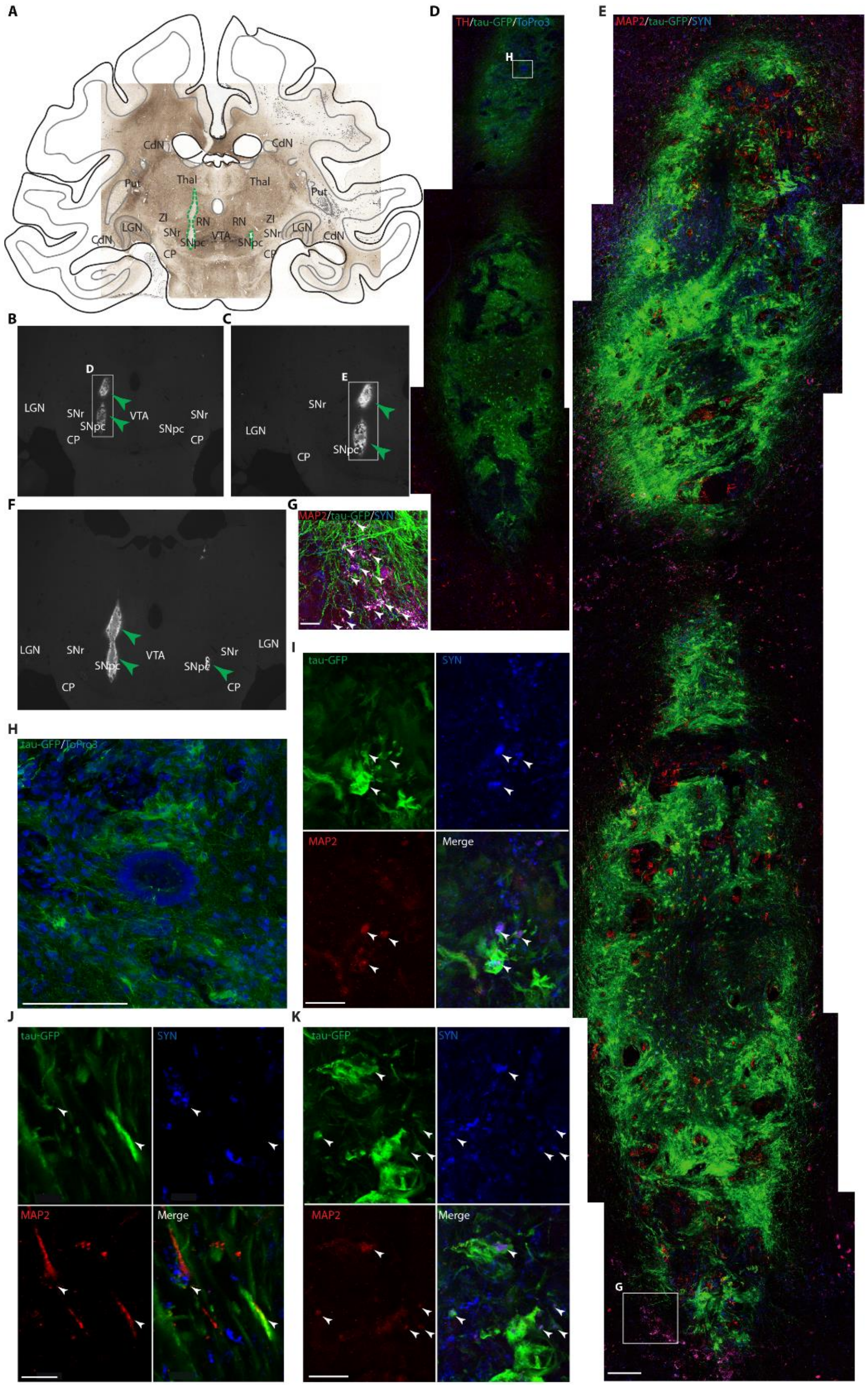


**Fig.S8: Grafted NPs in case 5.**

(**A**) Grayscale images from white-field microscope showing location of NPs in nigral transplant (green), drawings of the complete brain region from an adjacent section. (**B**), (**C**) and (**F**) Photomicrograph from fluorescent microscope acquisitions (tau-GFP$^+$ in white, filter 488nm, LeicaM165 FC). (**D**) and (**E**) Framed regions in, respectively (**B**) and (**C**). (**G**) Framed region in (**E**) showing accumulation of SYN$^+$ and MAP2$^+$ where tau-GFP$^+$ processes extend outside the core (white arrowheads. (**H**) Framed region in (**D**) showing host cell aggregate inside the graft-core. (**I**), (**J**), (**K**) Grafted cells (tau-GFP$^+$) co-localized with SYN$^+$ and MAP2$^+$. Scale bars: E 100µm; G 25µm; H 100 µm; I-K 5 µm.

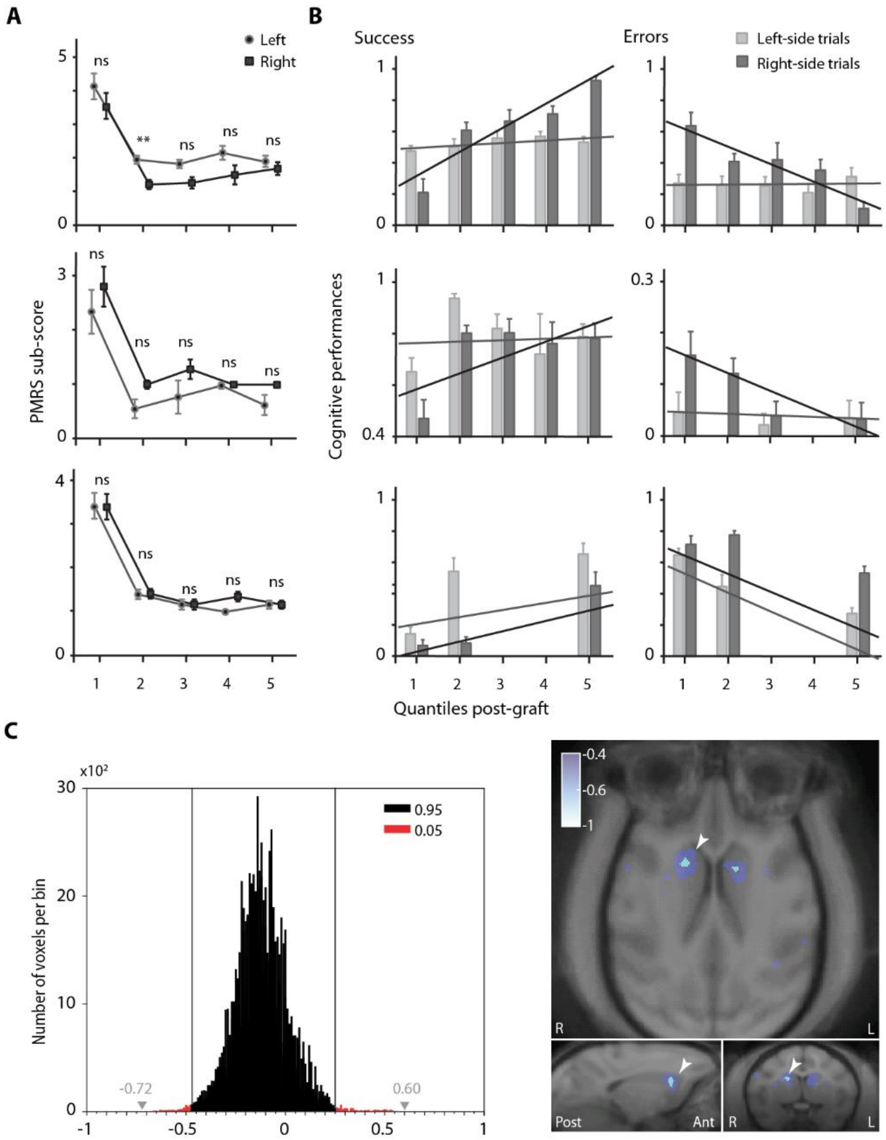


**Fig.S9: Hemispheric differences in graft-induced recovery.**

(**A**) Evolution of clinical sub-score in quantiles post-graft. Average sum of clinical score for resting tremor, ability to manipulate food, lower and upper limb movements for case1 (top), case5 (middle) and case4 (bottom) that was done separately scored for left (in grey) and right limbs (in black). Note that no clear difference could be related to NPs survival in left vs. right hemisphere. Case1 presented unilateral survival in posterior Put and bilateral survival in SN whereas case5 presented unilateral survival in SN and bilateral survival in posterior Put. Case4 is presented for completeness as no NPs were grafted in the Put (see **Table S3**). **p<0.001, ns p>alpha (Bonferroni corrected alpha=0.01, two-sample t-test assuming unequal variance). (**B**) Cognitive performances evolution in quantiles post-graft. Lines represent linear regression of average performances per quantile for left (grey lines) and right (black lines) trials separately. Recovery is characterized by an increase in Success and a decrease in Errors. Note the time-course asymmetry of recovery for cases 1 and 5 (top and middle row) and parallel time-course for case 4 (bottom row) when comparing left vs. right trials. Cases 1 and 5 presented unilateral survival whereas case4 presented bilateral survival in the anterior CdN (see **Table S3**). Mean±SE across quantiles. (**C**) The distribution of difference values of voxel-level $BP_{ND}$ Pre- vs. Post-graft, over all brain ROIs was used to calculate the $2.5^{th}$ and $97.5^{th}$ percentile boundaries (case1, vertical lines, -0.48 and +0.26 respectively). Grey arrowheads indicates min and max difference bins. Corresponding statistical thresholding of negative difference (right), note the minor but significant decrease in right anterior CdN (arrowheads). Positive difference shown in Fig.1C.

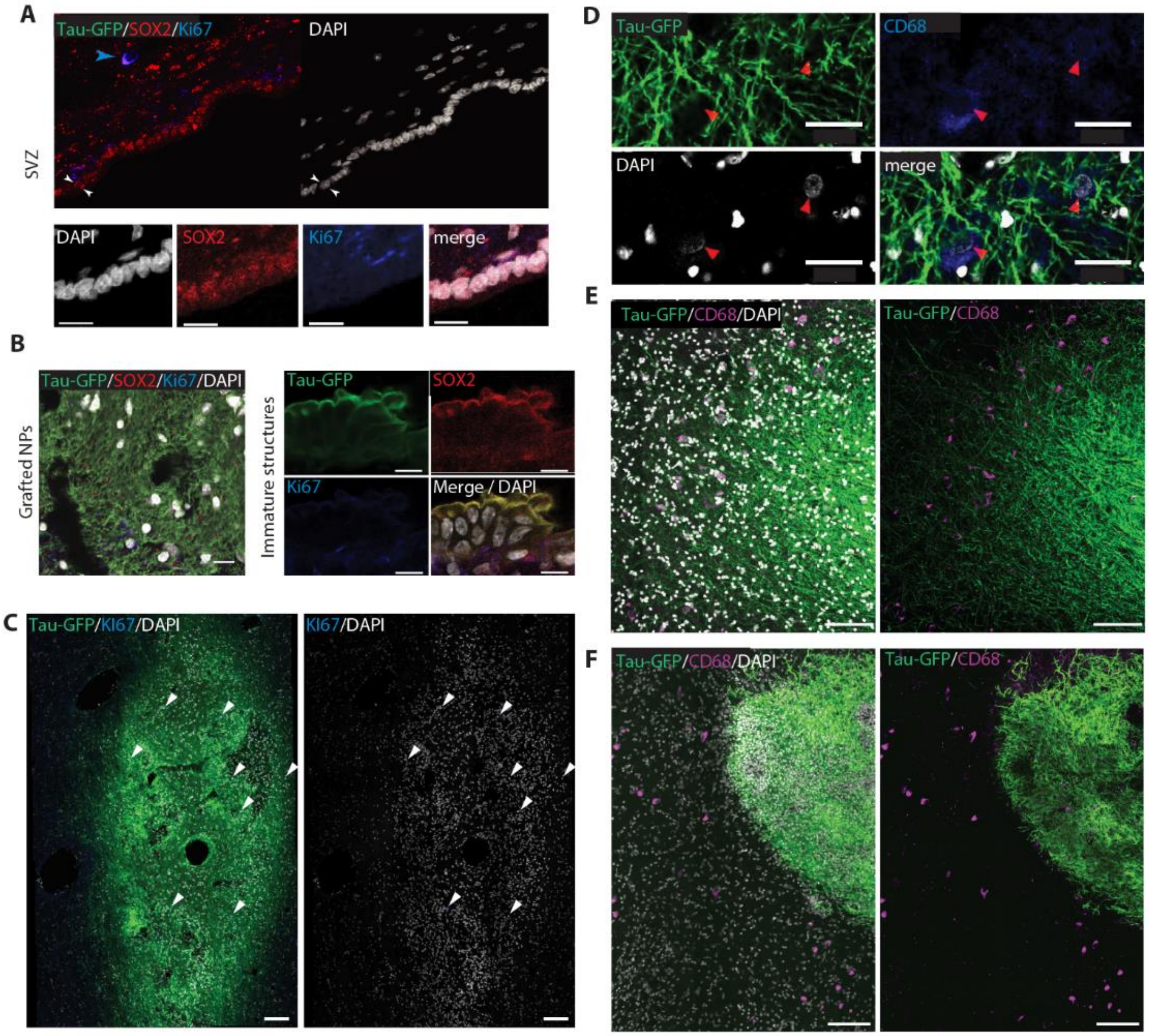


**Fig.S10: NPs show no overgrowth after transplantation and induce minor immune reaction.**

(**A**) Typical sub-ventricular zone section (SVZ, white arrowheads, case 5) displaying some proliferation (Ki67$^+$, blue arrowhead) and NP (SOX2$^+$) markers, scale bar 20µm. (**B**) Case5. Left, post-mortem immunostaining showing that no more NPs remain after graft (SOX2$^-$) from the tau-GFP population (in green) which neither shows sign of proliferation (Ki67$^-$); scale bar 13µm. Right, section at the level of a structure exhibiting an immature morphology which was however negative for SOX2$^-$ and Ki67$^-$; scale bar 10µm. (**C**) Few scattered Ki67$^+$/tau-GFP cells (white arrowheads) were observed in Case5, scale bar 100µm. (**D-F**) Few activated macrophages (**D**, red arrowheads, CD68$^+$) were present near grafted sites like presented here for SN graft sites in case 5 (**D** and **E**) and case 4 (**F**); scale bars (**D**) 30µm, (**E-F**) 100µm.

**Table S1: Longitudinal protocol design, repeated measures and case controls**

| Reference study | Protocol's Subparts | Neuroscience Disciplines | Present study cases | Total number of NHPs used |
|---|---|---|---|---|
| **Wianny et al.,2008** *Stem Cells (19)*<br><br>**Wianny et al.,2011** *Differentiation (15)* | ESCs line production and NPs derivation | Stem Cells Neurobiology | - | - |
| **Vezoli et al., 2011** *PLoS ONE (16)* | CTR; MPTP; Spontaneous Recovery | Neurobiology of Sleep and Circadian Rhythms; Cognitive Neurobiology; Clinical Neuroscience; Neurological Disease Models | 1, 2, 4 | 3 + monkey T |
| **Vezoli et al., 2011** *NeuroImage (17)* | CTR; MPTP; Spontaneous Recovery; Stable Symptoms | Neurobiology of Sleep and Circadian Rhythms; Cognitive Neurobiology; Clinical Neuroscience; Neuroimaging and Systems Neuroscience; Neuroanatomy; Neurological Disease Models | 1, 2, 3, 6 | 4 |
| **Fifel et al., 2014** *PLoS ONE (13)* | CTR; MPTP; Spontaneous Recovery; Stable Symptoms | Neurobiology of Sleep and Circadian Rhythms; Clinical Neuroscience; Neuroanatomy; Neurobiology of Disease | 1-6 | 6 + 5 monkeys (B, C, J, R, V) |
| **This study** | CTR; MPTP; Spontaneous Recovery; Stable Symptoms; Pre- vs. Post-Graft | Neurobiology of Sleep and Circadian Rhythms; Cognitive Neurobiology; Clinical Neuroscience; Neuroimaging; Anatomy and Histology; Neurobiology of Disease; Stem Cells Neurobiology; Translational Neuroscience | 1-6<br><br>3 additional control cases | 6 + 3 CTR |

Control procedures are detailed in the reference studies, identification of the cases used in the present study and the total number of animals used per study. Overall 15 different NHPs were involved in the longitudinal protocol design.

**Table S2: Total cumulated MPTP doses per case**

| | |
|---|---|
| **CASE 1** | **7.8 mg/kg (27 inj. SP and 12 inj. acute, 3months)** |
| **CASE 2** | **4.4 mg/kg (22 inj. SP/acute)** |
| **CASE 3** | **4.8 mg/kg (8 inj. SP and 16 inj. SP/acute, 5months)** |
| **CASE 4** | **4.4 mg/kg (11 inj. SP and 11 inj. acute, 26months)** |
| **CASE 5** | **3.0 mg/kg (15 inj. SP/acute)** |
| **CASE 6** | **2.4 mg/kg (12 inj. acute)** |

In cases 1, 3 and 4 MPTP intoxication was implemented in two phases with the first one consisting in chronic low-doses systemic injections (interval of at least 72-96H between injections (inj.), slowly progressive lesion (SP) that was halted in order to promote spontaneous recovery from motor symptoms. In all cases, MPTP intoxication ended after a period of acute low-doses injections (interval of 24h between injections). Time period between two MPTP intoxication periods indicated in months.

**Table S3. Cell survival following transplantation of NPs**

| | *Caudate nucleus* | | *Putamen* | | *Substantia nigra* | |
|---|---|---|---|---|---|---|
| | **LH** | **RH** | **LH** | **RH** | **LH** | **RH** |
| ***Case 1*** | $3.15\times10^{3}$ | 0 | $4.77\times10^{4}$ | 0 | $3.03\times10^{4}$ | $3.12\times10^{4}$ |
| **Case 2*** | - | - | - | - | - | - |
| ***Case 3*** | 0 | 0 | 0 | 0 | 0 | 0 |
| ***Case 4*** | $5\times10^{3}$ | $9\times10^{3}$ | NA | NA | $11\times10^{4}$ | 0 |
| ***Case 5*** | $5.67\times10^{4}$ | 0 | $2.52\times10^{4}$ | $4.95\times10^{3}$ | $18.4\times10^{4}$ | $<1\times10^{2}$ |
| ***Case 6*** | 0 | 0 | 0 | 0 | 0 | 0 |

Number of tau-GFP$^{+}$ cells estimated *post-mortem* after an average delay of 30±8 weeks post-graft. * Case2 had per-operative brain hemorrhage during NPs transplantation. Cases 1, 4 and 5 displayed functional recovery following cell-therapy (recovered group) whereas cases 3 and 6 did not (non-recovered group). Case 4 was not grafted with NPs in the Putamen.

**Table S4: Grafted cells number, survival rates, differentiation fates and comparison to literature**

| Grafted cells | Grafted cells number (per site) | Grafted sites | Survival rate (%) | Graft size ($mm^3$) | TH$^+$ | MAP2$^+$ | GFAP$^+$ | Graft type | Ref. |
|---|---|---|---|---|---|---|---|---|---|
| **NPs** | **$0.15\text{-}1\times10^6$** | **CdN, Put, SN** | **7±3** | **2.2±0.9** | **6±2%** | **16±4%** | **42±5%** | **Xeno** | **This study** |
| NSCs | $1\times10^6$ | CdN, SN | <10 | NA | <1% | 3% | NA | Xeno$^+$ | [56, 96] |
| | NA | CdN, Put | NA | 16-41 | <1.1% | 0% | 0% | Xeno$^+$ | [97] |
| | $10\times10^6$ | CdN, Put, SN | 10% | NA | 1.85% | NA | NA | Xeno$^+$ | [24, 98] |
| | $2\times10^6$ | CdN, SN | NA | NA | NA | NA | NA | Auto | [99] |
| | $2\times10^6$ | CdN, Put, SN | NA | NA | NA | NA | NA | Allo | [100] |
| DA neurons | $1.25\times10^6$ | CdN, Put | NA | NA | NA | NA | NA | Xeno$^+$ | [101] |
| | $2.4\times10^6$ | Put | NA | NA | <0.75% | 39% | NA | Xeno$^+$ | [102] |
| | $1\times10^6$ | CdN, Put | NA | NA | 10% | NA | NA | Xeno$^+$ | [103] |
| | $2.5\times10^6$ | Put | NA | 5.4-14 | <0.1% | NA | NA | Allo | [104] |
| | $0.15\text{-}0.3\times10^6$ | Put | 1.3-2.7 | NA | 0.4% | NA | NA | Allo$^+$ | [105] |
| | $0.25\text{-}0.75\times10^6$ | CdN, Put, SN | <1.5 | NA | NA | 63% | 22% | Auto | [22] |
| | $5\times10^6$ | Put | NA | NA | <0.1% | NA | NA | Auto | [106, 107] |
| | NA | CdN, Put | NA | NA | 0.4% | NA | NA | Auto | [108] |
| | $2.4\times10^6$ | Put | NA | 39.4±21 | 33±24% | NA | NA | Xeno$^+$ | [109] |
| | $0.5\text{-}7.3\times10^6$ | CdN, Put ± SN | NA | 0.63-36.5 | * | NA | NA | Auto/Allo | [73] |
| | $2\times10^6$ | CdN, Put | 6-53% | ~100-150 | 5.2-8.1% | NA | NA | Xeno | [83] |
| Other | $0.3\text{-}0.5\times10^6$ | CdN or Put | NA | NA | NA | NA | NA | Xeno | [110] |
| | $4\times10^6$ | CdN | NA | NA | NA | NA | NA | Allo | [111] |
| | $0.5\text{-}3\times10^6$ | CdN, Put, SN | NA | NA | NA | NA | NA | Allo | [112] |
| | NA | Put | NA | NA | ** | NA | NA | Allo | [113] |
| | $0.2\text{-}0.4\times10^6$ | CdN, Put, SN | 10-50 | NA | NA | NA | NA | Allo | [114] |

*$0.6\pm0.7\times10^4$ (Allo) and $9.1\pm5.8\times10^4$ (Auto) TH$^+$ cells at grafted location; **80-100 TH$^+$ cells at grafted location; $^+$immunosuppressive treatment. Min-Max and Mean±SE.

**Table S5: Clinical motor, cognitive and functional impact of grafts described in Table S4**

| Grafted cells | Clinical motor score | Cognitive behavior | Circadian rhythm | F-DOPA | DAT | Other, comments | Ref. |
|---|---|---|---|---|---|---|---|
| NPs | Improvement | Improvement | No effect | Increase | Increase | - | This study |
| NSCs | Improvement | NA | NA | NA | NA | - | [56, 96] |
| | NA | NA | NA | NA | NA | Asymptomatic animals | [97] |
| | No effect | NA | NA | NA | NA | Increase in striatal DA | [24, 98] |
| | Improvement | NA | NA | NA | Increase | No quantification | [99] |
| | Improvement | NA | NA | NA | Increase | No quantification | [100] |
| DA neurons | NA | NA | NA | NA | NA | - | [101] |
| | Improvement | NA | NA | Increase | NA | - | [102] |
| | NA | NA | NA | NA | NA | - | [103] |
| | Improvement | NA | NA | Increase | NA | - | [104] |
| | Improvement | NA | NA | NA | NA | - | [105] |
| | NA | NA | NA | NA | NA | - | [22] |
| | NA | NA | NA | NA | NA | - | [106, 107] |
| | Improvement | NA | NA | NA | NA | - | [108] |
| | Improvement | NA | NA | NA | NA | - | [109] |
| | Improvement (Autograft) | Reduced depressive signs | NA | NA | NA | Increase of VMAT2 (Autograft only) | [73] |
| | Improvement | NA | NA | NA | NA | increase in striatal DA (HPLC) | [83] |
| Other | Improvement | NA | NA | NA | NA | Change in FDG patterns | [110] |
| | Asymptomatic animals | NA | NA | NA | NA | Increase in striatal DA metabolite | [111] |
| | NA | NA | NA | NA | Increase | Graft before MPTP, increase in striatal DA, SERT | [112] |
| | Improvement | NA | NA | Increase | NA | - | [113] |
| | Improvement | NA | NA | NA | NA | - | [114] |

SPECT: Single-photon emission computed tomography; FDG: fluorodeoxyglucose; VMAT2: Vesicular MonoAmine Transporter type 2; HPLC: High-Performance Liquid Chromatography.

#### Table S6: Antibodies list

| Name | Supplier | Reference |
|---|---|---|
| Rabbit anti-GFAP | Dako | Z0334 |
| Chicken anti-GFP | Life Technologies | A10262 |
| Mouse anti-MAP2 | SIGMA | M4403 |
| Mouse anti-NESTIN | Millipore | MAB5326 |
| Mouse Tuj1 | SIGMA | T8660 |
| Goat anti-SOX2 | Santa Cruz | sc-17320 |
| Rabbit anti-Ki67 | Thermo scientific | RM-9106 |
| Goat anti-GDNF | R&D systems | AF-212-NA* |
| Mouse anti-CD68 | R&D systems | MAB 20401 |
| Rabbit anti-SYNUCLEIN | ProteinTech | 17785-1-AP |
| Rabbit anti-TYROSINE HYDROXYLASE | Chemicon | VPA151 |
| Mouse anti- TYROSINE HYDROXYLASE | Millipore | MAB318 |
| Rat anti-DOPAMINE TRANSPORTER (DAT) | Millipore | MAB369 |
| Rabbit anti-SEROTONIN TRANSPORTER (SERT) | Millipore | AB9322 |
| Goat anti-Chicken alexa 488 | Life Technologies | A11039 |
| Goat anti-Mouse alexa 647 | Life Technologies | A21235 |
| Goat anti-Rabbit alexa 555 | Life Technologies | A21428 |
| Donkey anti-Goat alexa 555 | Life Technologies | A21432 |
| Donkey anti-Rabbit alexa 647 | Life Technologies | A31373 |
| Donkey anti-Rabbit alexa 555 | Life Technologies | A31572 |
| Donkey anti-Mouse alexa 647 | Life Technologies | A31571 |
| Donkey anti-Chicken dylight 488 | Jackson | 703-485-155 |

*not shown in results

#### Table S7: Primers for semi-quantitative RT-PCR

| Gene | Forward Primer (5′-3′) | Reverse Primer (5′-3′) |
|---|---|---|
| **β actin** | AAACTGGAACGGTGAAGGTG | TCAAGTTGGGGGACAAAAAGG |
| **AhD2** | GTTGTCAAACCAGCAGAGCA | CAAGTCGGCATCAGCTAACA |
| **DRD2** | GGAGGTGGTAGGTGAGTGGA | GGAGATGGTGAAGGACAGGA |
| **EN1** | CTAGCCAAACCGCTTACGAC | GCAGAACAGACAGACCGACA |
| **GAD67** | ATTCTTGAAGCCAAACAG | TAGCTTTTCCCGTCGTTG |
| **GFAP** | GGCAGGGCATGACTTTGTTC | TAAGAAGGGACCGCAAGAGG |
| **LMX1A** | CCTGCAGGAAGGTGAGAGAG | GTCGTCGCTATCCAGGTCAT |
| **LMX1B** | GCAGCGGCTGCATGGAGAAGATCGC | GGTTCTGAAACCAGACCTGGACAAC |
| **MAP2** | CAGCAAAGGGATACTTTCAC | ATGCTTTTTGTTGCTTCTTC |
| **MUSASHI** | CGAGCTCGACTCCAAAACAATT | TCTACACGGAATTCGGGGAACT |
| **NESTIN** | CGTCTTGGATCTTTGCTCCC | GGGCTCTGATCTCTGCATCT |
| **NGN2** | CCGAGACCTTGGAGTTGAAG | CGTTTGCAATCGTGTACCAG |
| **NURR1** | CTCCCAGAGGGAACTGCACTTCG | CTCTGGAGTTAAGAAATCGGAGCTG |
| **OCT4** | CGACCATCTGCCGCTTTGAG | CCCCCTGTCCCCCATTCCTA |
| **PAX2** | TGTGTCAGCAAAATCCTGGGCAGGT | TGCTGAACTTTGGTCCGGATGAT |
| **PAX6** | CATGCAGAACAGTCACAGCGG | CCCATCTGTTGCTTTTCGCTA |
| **SOX2** | CCCCCGGCGGCAATAGCA | TCGGCGCCGGGGAGATACAT |
| **TH** | ATGGCTGAGCCCCGCCAGGAG | TCACAAACCCTGCTTGGCC |
| **c-RET** | CGACCTCATCTCATTTGCC | AATCTTCATCTTCCGCCCC |

#### Table S8: Group-level stats of Fig.1B

| | | PMRS | ORDT success | ORDT errors | LD ratio |
|---|---|---|---|---|---|
| CTR | | ns (score = 0) | ns | ns | t = 4.6, p = 1.8E-04, n = 128 |
| MPTP | Q1 | ns | ns | ns | t = 0.2, p = 1.5E-02, n = 42 |
| | Q2 | t = 0.6, p = 3.4E-02, n = 38 | ns | ns | ns |
| | Q3 | t = 3.4, p = 5.6E-04, n = 39 | ns | ns | ns |
| | Q4 | ns | ns | ns | ns |
| | Q5 | ns | ns | ns | ns |
| Post-MPTP | Q1 | ns | ns | ns | ns |
| | Q2 | t = -9.4, p = 6.8E-19, n = 79 | ns | ns | ns |
| | Q3 | t = -3.5, p = 2.4E-02, n = 68 | ns | ns | ns |
| | Q4 | ns | NA | NA | ns |
| | Q5 | ns | NA | NA | ns |
| Post-graft | Q1 | t = -7.4, p = 7.8E-12, n = 164 | ns | ns | ns |
| | Q2 | t = -15.4, p = 7.3E-49, n = 189 | ns | ns | ns |
| | Q3 | t = -15.9, p = 1.6E-51, n = 192 | t = 3.3, p = 3.6E-02, n = 36 | t = -5.3, p = 6.2E-06, n = 36 | ns |
| | Q4 | t = -17.3, p = 1.5E-60, n = 195 | t = 5.4, p = 4.9E-06, n = 26 | t = -4.8, p = 9.1E-05, n = 26 | t = 4.4, p = 4.3E-04, n = 175 |
| | Q5 | t = -15.5, p = 3.3E-49, n = 153 | t = 5.1, p = 2.3E-05, n = 36 | t = -4.7, p = 1.2E-04, n = 36 | t = 9.1, p = 1.1E-17, n = 121 |

One-way ANOVA *recovered* vs. *non-recovered*, adjusted p-values with Bonferroni correction for multiple comparisons.
PMRS: Parkinsonian Monkey Rating Scale – Clinical motor score; ORDT: object retrieval detour task – Cognitive performances;
LD ratio: Light/Dark ratio – Chronobiological rhythms.
